# Digital-robotic markers for hallucinations in Parkinson’s disease

**DOI:** 10.1101/2023.06.14.544929

**Authors:** Louis Albert, Jevita Potheegadoo, Bruno Herbelin, Fosco Bernasconi, Olaf Blanke

## Abstract

Hallucinations are frequent non-motor symptoms in Parkinson’s disease (PD) associated with dementia and higher mortality. Despite their high clinical relevance, current assessments of hallucinations are based on verbal self-reports and interviews that are limited by important biases. Here, we used virtual reality (VR), robotics, and digital online technology to quantify presence hallucination (PH) in laboratory and home-based settings. We establish that elevated numerosity estimation of digital humans in VR is a digital marker for experimentally induced PH in healthy participants, as confirmed across several control conditions and analyses. We translated the digital marker (numerosity estimation) to an online procedure that 170 PD patients carried out remotely at their homes, revealing that PD patients with disease-related PH (but not control PD patients) showed higher numerosity estimation. Numerosity estimation enables quantitative monitoring of hallucinations, is an easy-to-use unobtrusive online method, reaching people far away from medical centers, translating neuroscientific findings using robotics and VR, to patients’ homes without specific equipment or trained staff.

## Introduction

Parkinson’s disease (PD) is the second most common neurodegenerative disease following Alzheimer’s disease, affecting approximately 3% of the population over 65 years of age (Kalia and Lang 2015). Although PD is defined primarily as a movement disorder (i.e. resting tremor, rigidity, bradykinesia), it is a heterogeneous disorder, also affecting several non-motor systems and manifesting in a wide variety of non-motor symptoms, including hallucinations (Postuma and Berg 2016). Hallucinations in PD are highly prevalent (with approximately half of PD patients experiencing hallucinations) and can reach up to 70% during later stages of the disease (Ffytche and Aarsland 2017). Critically, hallucinations have been associated with a more severe form of PD with negative clinical outcomes including dementia, depression, early home placement, and a higher mortality (ffytche and Aarsland 2017; Diederich et al. 2009; Fénelon et al. 2000; Marinus et al. 2018; Kataoka and Ueno 2015; Bernasconi, Pagonabarraga, et al. in press; Lenka et al. 2019; Forsaa et al. 2010).

Hallucinations in PD have been categorized into formed (well-structured or complex) visual hallucinations and so-called minor hallucinations, which include presence hallucinations (PH) and passage hallucinations, and visual illusions (Fénelon et al. 2000; Ravina et al. 2007). Visual hallucinations generally occur at the middle to late stage of the disease, and several studies have identified visual hallucinations as a risk factor for more rapid cognitive decline and dementia in PD (Aarsland et al. 2003; Anang et al. 2014; Galvin, Pollack, and Morris 2006; Uc et al. 2009). However, because visual hallucinations occur at a more advanced stage of the disease, with cognitive decline already present, they are not suitable as an early marker of cognitive decline in PD. This differs for minor hallucinations, which are usually experienced at earlier stages of the disease (Ffytche and Aarsland 2017; Lenka et al. 2019), and can even precede parkinsonian motor symptoms, testifying to the importance to include them in detailed clinical evaluations (Pagonabarraga et al. 2016). Recent data show that minor hallucinations are not only the earliest hallucinations occurring in PD, but that they also share brain alterations with visual hallucinations (Pagonabarraga et al. 2014; H. Bejr-kasem et al. 2019) and are linked to more rapidly developing cognitive deficits (Bernasconi, Pagonabarraga, et al. in press; H. Bejr-kasem et al. 2021; Bernasconi et al. 2021), underlining their potential role as an early marker for dementia (Pagonabarraga et al. 2014; H. Bejr-kasem et al. 2021; Bernasconi et al. 2021; Bernasconi, Pagonabarraga, et al. in press).

Despite their clinical relevance, the diagnosis and investigation of hallucinations is hampered by difficulties to examine them in real-time and quantify them reliably. Hallucinations are subjective-private experiences (Rogers, Keogh, and Pearson 2021) and self-reports are inadequate for precisely representing and quantifying hallucinations (Bernasconi, Blondiaux, et al. 2022; Rogers, Keogh, and Pearson 2021). Yet, the current gold standard in medical practice for assessing type, frequency, and intensity of hallucinations is based on verbal self-reports and interviews of patients and interpretations by clinicians. However, verbal reports only capture some aspects of conscious experience and are prone to many biases (i.e., we dispose of a limited ability to accurately recall and report past conscious experiences, with an unknown reliability; many aspects of these conscious experiences cannot be accurately quantified in verbal reports; verbal reports are prone to the interviewer biases (Rogers, Keogh, and Pearson 2021; Nisbett and Wilson 1977)). These limitations are further exacerbated, because for many patients reporting their hallucinations may be affected by lack of insight and/or fear of stigmatization, which often refrains patients from reporting them (Fénelon et al. 2011).

New procedures and methods have been developed to overcome some of these limitations, allowing a quantitative assessment of hallucinations and to investigate and induce hallucinations in controlled laboratory setting, in both healthy and clinical populations (i.e., Bernasconi, Blondiaux, et al. 2022). Of relevance for PD, a sensorimotor robotic procedure has been shown to induce repeatedly a specific and clinically relevant hallucination, presence hallucination (PH; the vivid sensation that another person is nearby when no one is actually present and can neither be seen nor heard (Arzy et al. 2006)), in healthy participants and in PD patients (Blanke et al. 2014; Bernasconi, Blondiaux, et al. 2022). Such real-time induction of a clinically relevant hallucination in healthy participants has permitted to identify the brain mechanisms underlying these aberrant perceptions without the confounds present in clinical populations (i.e., co-morbidities not related to hallucinations) (Bernasconi et al. 2021; Bernasconi, Blondiaux, et al. 2022; Rogers, Keogh, and Pearson 2021). Moreover, the translation of the procedure and methods to patients with PD confirmed that robotically induced PH are clinically relevant, because they shared key phenomenological aspects with PD patients’ spontaneous PH in daily life and PD patients with spontaneous PH were more sensitive to the robotically induced PH procedure (Bernasconi, Blondiaux, et al. 2022). However, while the robotics-based approach enabled the investigation of PH in real-time within a controlled environment (i.e., overcoming several limitations of earlier hallucination research), the procedure still relied on explicit ratings (e.g., questionnaires), which are sensitive to participant and experimenter biases (Rogers, Keogh, and Pearson 2021; Nisbett and Wilson 1977; Wegner 2003), but may be overcome by implicit behavioral proxies (*APA Dictionary of Psychology* 2007), as applied to the sense of self-location (Lenggenhager, Mouthon, and Blanke 2009; Nakul et al. 2020), spatial thought (Parsons 1987; Shepard and Metzler 1971), or agency judgements (Moore and Obhi 2012; Haggard, Clark, and Kalogeras 2002)

Here, we designed a novel and fully controlled behavioral numerosity task with visual digital humans (and with visual control objects: control task), using immersive Virtual Reality (VR) technology that we combined with our robotic PH-induction method (Bernasconi et al. 2021) (study 1). We combined VR with the robotic system and determined whether the new task is an implicit, quantitative and behavioral marker for PH. Our findings reveal an overestimation bias for human stimuli that is robust and based on many repeated trials, which is observed in the PH-inducing condition, and absent in the control task (object condition), establishing the overestimation of digital humans as an implicit behavioral marker for PH. Based on these results and our previous finding that PD patients with minor hallucinations show heightened sensitivity to robotically induced PH (Bernasconi et al. 2021), we next developed a home-based online numerosity task with digital humans and tested a large group of PD patients at their home (study 2). Online data reveal a larger overestimation bias for digital humans in PD patients with PH (PD-PH) as compared to control PD patients without hallucinations (PD-nH), demonstrating an implicit digital online marker for early hallucinations in PD.

## Results

### Experiencing an invisible presence increases visual overestimation bias for numerosity estimations of visual digital humans (study 1)

#### Estimation of human stimuli using immersive VR

To test whether classical NE effects observed for different visual stimuli such as dots (Arrighi, Togoli, and Burr 2014; Cicchini, Anobile, and Burr 2014; Fornaciai and Park 2020; Jevons 1871; Kaufman et al. 1949; Poulton 1979; Minturn and Reese 1951; Krueger 1984), squares (Anobile et al. 2022; O’Hearn, Hoffman, and Landau 2011; Anobile et al. 2020) or cartoon animals (Leibovich-Raveh et al. 2018) can also be observed for more complex and ecologically valid stimuli such as digital humans in a room (as shown in a VR scenario), we developed a new immersive 3D VR paradigm. To maximize immersion and strengthen the ecological validity of our experiment, healthy participants were first immersed in a virtual reconstruction of the actual testing room, including the participants’ actual location and orientation in the exact same room of our laboratory. Stimuli consisted in brief displays of the 3D VR environment, which contained a varying number of digital humans (Figure 1; Extended Data Figure 1; Figure S1; Video S1) that were equally positioned in the virtual room, in front of the participant, at least at a distance of 1.75 virtual meters (vm), and within the near peripheral field of view. The 3D VR scene was displayed to participants on a Head-Mounted Display (HMD) (Oculus Rift CV1). Participants were asked to indicate the number of humans (NEH task) they perceived in the room, as fast and as precise as possible. A control condition was also performed (number of objects (NEO) task; Figure 1; Figure S1; Video S2).

**Figure 1.**
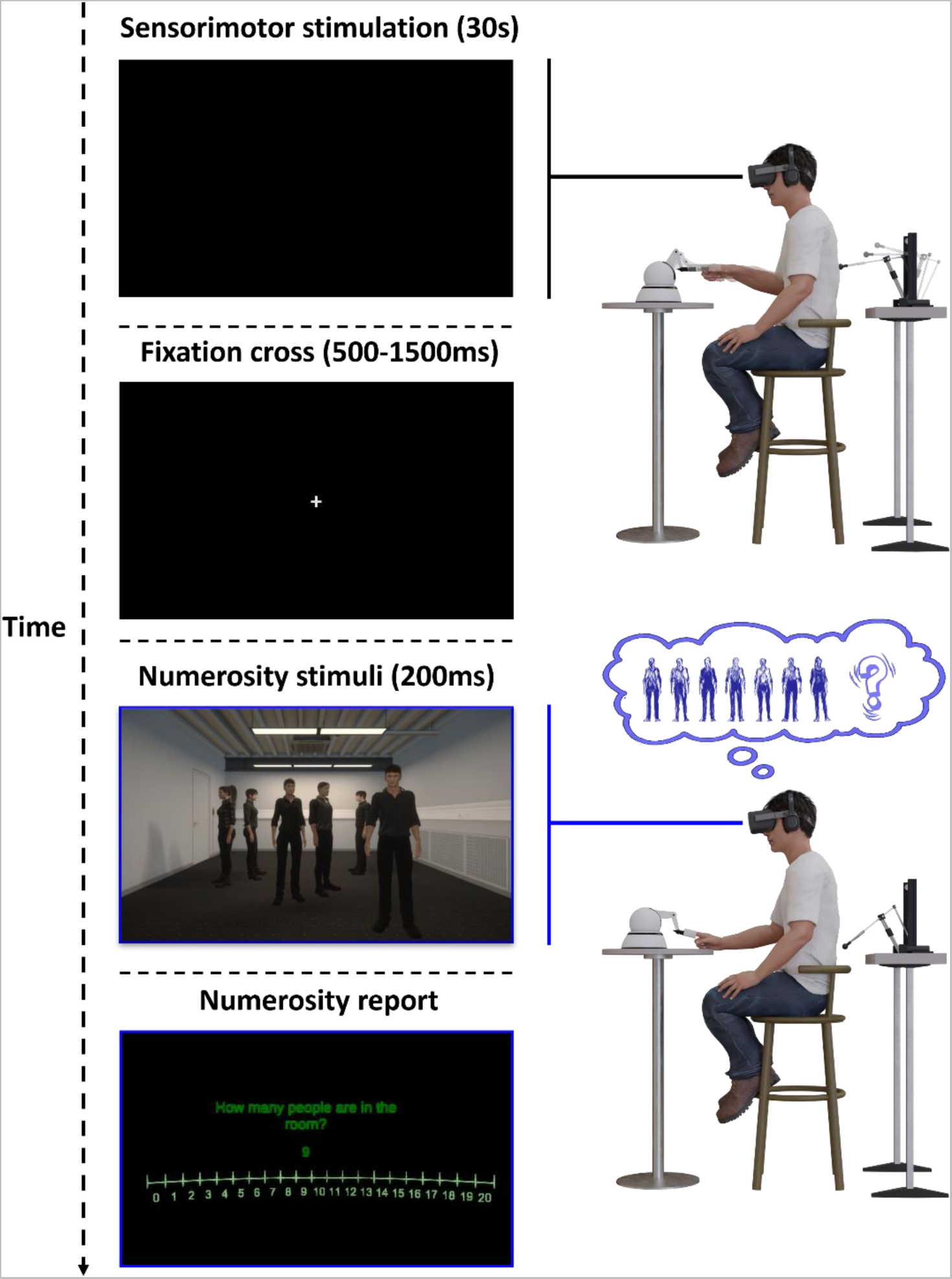
Integrating sensorimotor robotic stimulation, virtual reality, and numerosity estimation task (study 1). In each NEH trial, participants first manipulated the robotic system for 30 seconds (either in the asynchronous (500ms delay; PH-inducing condition) or the synchronous condition (0ms delay)). This was followed by the appearance of a fixation cross (500-1500ms), indicating to participants that they could stop moving the robotic system. Then, a scene containing a different number of people (range 5-8) was shown for 200ms and participants had to estimate the number of people they saw. All visual stimuli were shown in immersive virtual reality on a head-mounted display (see methods for further detail).

To determine the lower bound of the estimation range of our NEH stimuli and select the range of presented numerosities in study 1, we conducted an online pilot study (see Note S1) in 28 healthy participants. In this online preliminary study, participants were asked to indicate the number of humans they perceived in flashed human stimuli over a broad range of numerosities (ranging from 1 to 24). This online preliminary study indicated the lower bound of the estimation range of our NEH stimuli to be 5. For all methodological aspects and detailed results of this online pilot study see Note S1.

First, as predicted based on the previous literature (Jevons 1871; Kaufman et al. 1949; Minturn and Reese 1951), and in agreement with our preregistered hypothesis (Albert et al. 2021), our data show that NE is modulated by the number of stimuli (humans or objects) presented in the virtual room (i.e., presented numerosity; F(3, 2197)=1946; p<0.001; main effect; Table S3). In particular, we observed that NE increases with the number of presented numerosities, independently of the type of stimulus (Table S2; Table S3). Second, and in agreement with our hypothesis (Albert et al. 2021), additional post-hoc analysis showed that participants mean NE is significantly higher than the visually presented numerosity for the range of presented numerosity (5 to 8) (Figure 2; Table S2). This behavior is typical and has been observed with dots stimuli for numerosities just above the subitizing range (Fornaciai and Park 2020; Jevons 1871; Minturn and Reese 1951). These data confirm and extend two well-known effects from classical NE (carried out with dots on 2D computer screens: numerosity main effect, overestimation above the subitizing range) (Fornaciai and Park 2020; Jevons 1871; Minturn and Reese 1951), that we also observed in our pilot online study (note S1). We here report them for the first time for the numerosity estimation of humans and control objects in immersive VR using a HMD.

**Figure 2.**
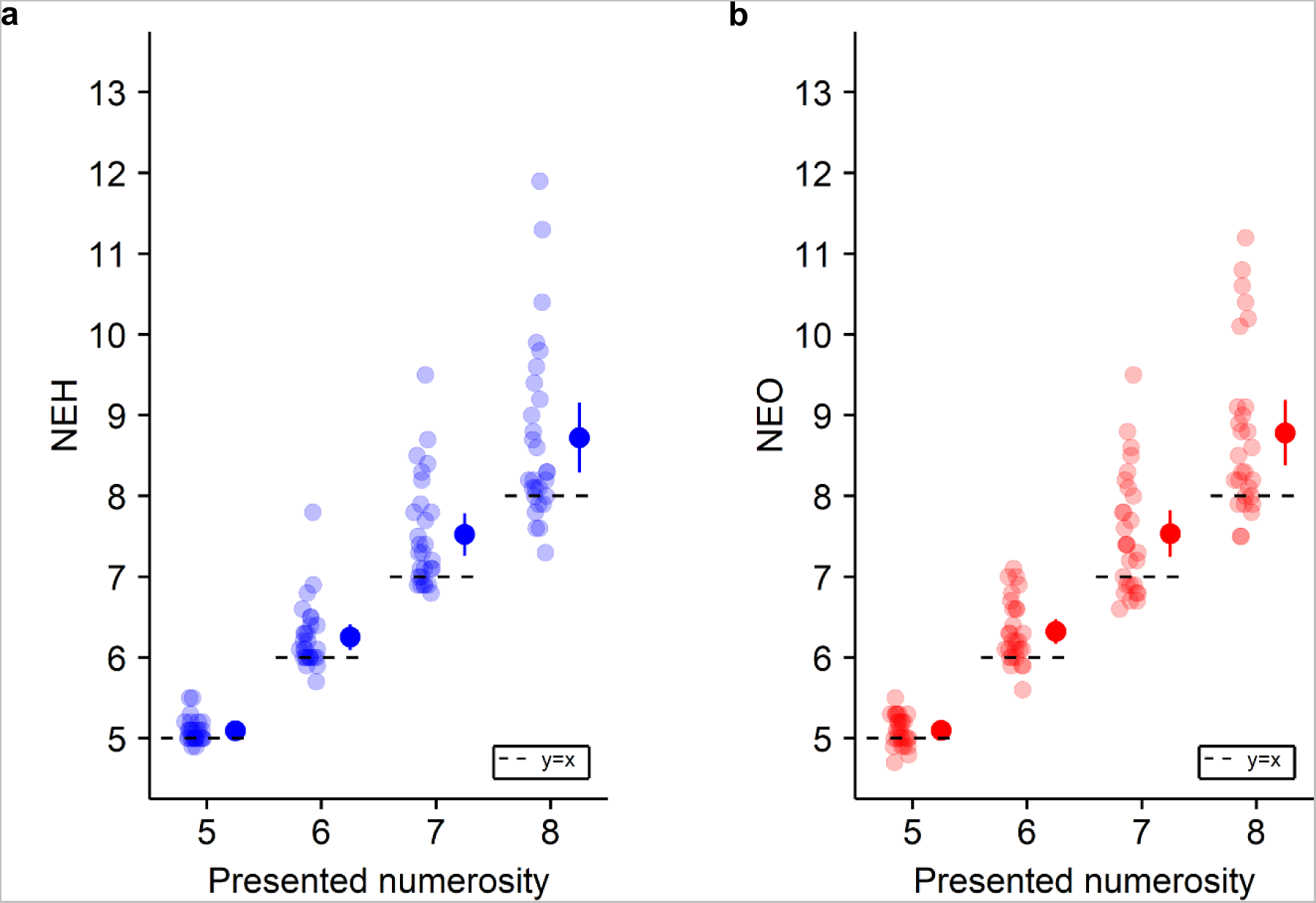
Numerosity estimation (study 1). General numerosity estimation performance for each tested numerosity in the (a) NEH and (b) the NEO task (study 1). Each dot indicates the individual NEH mean (over the trials) estimate for the corresponding presented numerosity. The dots with the bar on the right sides indicate the in-between subject mean for each presented numerosity. Note the general overestimation bias in NEH and NEO. Error bar represents 95% confidence interval.

#### Higher overestimation in NEH is associated with PH

To assess whether the magnitude of the NEH overestimation is a valid implicit measure for robot-induced PH, our NEH stimuli and procedure were integrated with the robotic sensorimotor paradigm (Bernasconi et al., 2022) that has been shown previously to induce robot-induced PH in healthy participants (Bernasconi et al. 2021; Blanke et al. 2014; Orepic et al. 2021; Serino et al. 2021; Salomon et al. 2020). To induce robot-induced PH, participants were asked to perform repetitive movements to operate a robot placed in front of them, which was combined with a back robot providing tactile feedback to the participants’ backs with a delay of 500ms (asynchronous sensorimotor stimulation) (Figure 1). A second sensorimotor condition (synchronous sensorimotor stimulation) served as a control condition (participants performed the same repetitive movements to operate the front robot and received the same tactile feedback on their backs, and with the same spatial conflict, but without the additional 500ms temporal delay of the asynchronous condition). On a trial-by-trial basis, participants performed NEH (using the immersive VR procedure) immediately after each sensorimotor stimulation phase of 30 seconds (i.e., either asynchronous or synchronous condition), for a total of 40 NEH trials (for protocol see Figure 1 and Extended Data Figure 1). To reinforce the context of our NEH task, as a habituation phase, prior to each task, participants were immersed in the virtual room for one minute. During this habituation phase several digital humans moved and were seen discussing among themselves in the virtual environment. We tested whether: i) sensorimotor stimulation is associated with changes in NEH and, ii) in particular, if the magnitude of NEH overestimation can be used as an implicit marker of PH in the asynchronous versus synchronous sensorimotor condition. That is, we hypothesize that the robot-induced invisible PH would increase the NE bias of participants when exposed to the digital humans displayed on the HMD. Critically, we predicted that this effect should be larger in the asynchronous versus synchronous condition and it would be absent in our control NEO task (where instead of digital humans, digital control objects are shown in the same virtual room, at the same positions and orientations, and for the same numerosities) (Extended Data Figure 1; Figure S1). In agreement with our preregistered hypothesis (Albert et al. 2021), we observed an overestimation in the PH-inducing asynchronous condition that was specific to NEH (i.e., absent for NEO). Indeed, our results show that NE is significantly (F(1, 2197)=11.5; p<0.001; Interaction; Table S3) modulated by the synchrony of sensorimotor stimulation and the type of stimuli (NEH vs. NEO). Critically, and in agreement with our preregistered hypothesis (Albert et al. 2021), post-hoc comparisons showed that sensorimotor robotic stimulation significantly modulates NEH (t(2197)=−2.9; p=0.003; Table S3) (Figure 3a; Figure 3b) and that the PH-inducing asynchronous condition induces a stronger NEH overestimation bias than the synchronous control condition. Moreover, this effect was only present when estimating the number of humans in the virtual room, as it was absent for NEO (t(2197)=1.87; p=0.06; Table S3) (Figure 3c; Figure S5). Collectively, these data show that the NEH overestimation depends on sensorimotor stimulation, that it is larger in the asynchronous condition versus synchronous control condition and that this modulation is not observed for the NEO task. These findings reveal an overestimation bias for human stimuli in the PH-inducing asynchronous condition, suggesting that NEH is an implicit behavioral marker or proxy for PH.

**Figure 3.**
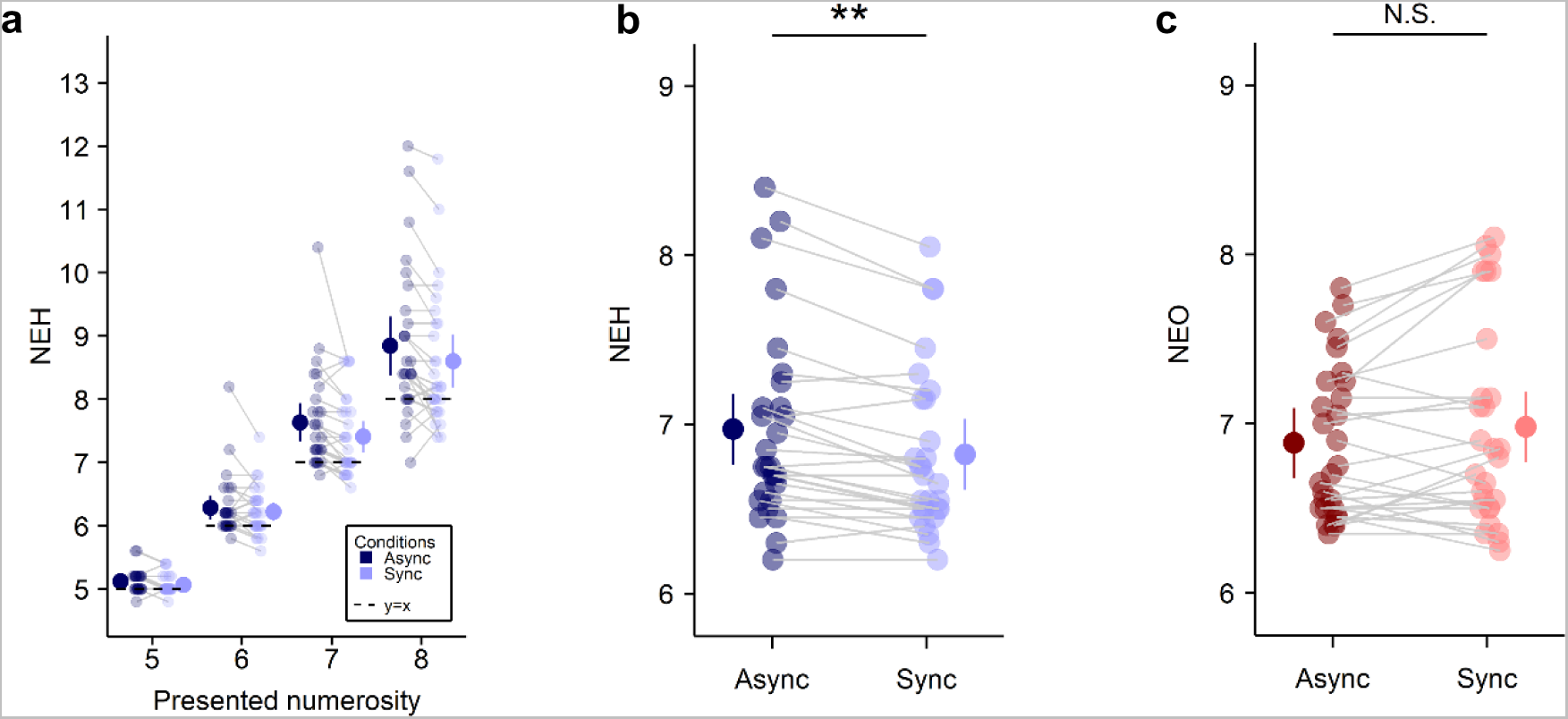
NEH and NEO as a function of sensorimotor stimulation (synchronous, asynchronous) (study 1). (a) Task performance is shown for each presented numerosity in the NEH task, separately for the asynchronous and synchronous condition. Each linked pair of dots indicates the individual NEH mean estimate for the corresponding numerosity in asynchronous condition (dark blue) and synchronous condition (light blue). The dots with the bar on the left and right sides indicate the mixed-effects linear regression between asynchronous (dark blue) and synchronous (light blue) sensorimotor stimulation for each tested numerosity. (b) NEH (asynchronous versus synchronous condition). Each linked pair of dots indicates the individual NEH mean estimate in asynchronous condition (dark blue) and synchronous condition (light blue). The dots with the bar on the left and right sides indicate the mixed-effects linear regression between asynchronous (dark blue) and synchronous (light blue) sensorimotor stimulation. (c) NEO (asynchronous versus synchronous). Each linked pair of dots indicates the individual NEO mean estimate in asynchronous condition (dark red) and synchronous condition (light red). The dots with the bar on the left and right sides indicate the mixed-effects linear regression between asynchronous (dark red) and synchronous (light red) sensorimotor stimulation. Error bar represents 95% confidence interval. **P ≤ 0.01. N.S., not significant.

Additional analysis ensured that the observed differences between NEH and NEO are not explained by differences in task difficulty. Thus, we observed no significant differences in response times between NEH and NEO (i.e., type of stimuli; F(1,2197)=0.73; p=0.39; no main effect plus no interactions; Table S4) (Figure S3; Figure S4; Figure S6; Figure S7). This analysis also indicated no effect of robotic sensorimotor stimulation on response times (i.e., type of robotic sensorimotor stimulation; F(1,2197)=3.01; p=0.08; no main effect, no interaction; Table S4), suggesting that the different robotic sensorimotor stimulation conditions did not affect NE task difficulty nor alertness. This is supported by the fact that the NE task is not performed during but just after the robotic stimulation. These results consolidate our findings that the overestimation observed in the asynchronous versus synchronous condition and in NEH versus NEO cannot be explained by differences in task difficulty.

#### PH is stronger in asynchronous versus synchronous condition

To ensure the successful induction of PH during the robotic sensorimotor stimulation, we additionally administered, at the beginning of the experiment and prior to the NE task, a previously used comprehensive questionnaire about PH (Table S1, Figure S2). In line with previous results observed with the robotic sensorimotor paradigm (Bernasconi et al. 2021; Blanke et al. 2014; Orepic et al. 2021; Serino et al. 2021; Salomon et al. 2020), the present participants reported higher PH in the asynchronous sensorimotor condition (mean = 2.29, SD = 1.96) compared to the synchronous condition (mean = 1.50, SD = 1.99) (χ^2^ (1, N = 28) = 12.00, p < 0.001) (Figure 4a; Figure S2; Table S1). Other robot-induced bodily experiences (illusory self-touch, passivity experience and loss of agency) were also compatible with previous findings (Bernasconi et al. 2021; Blanke et al. 2014; Serino et al. 2021; Salomon et al. 2020) (Table S1, Figure S2).

**Figure 4.**
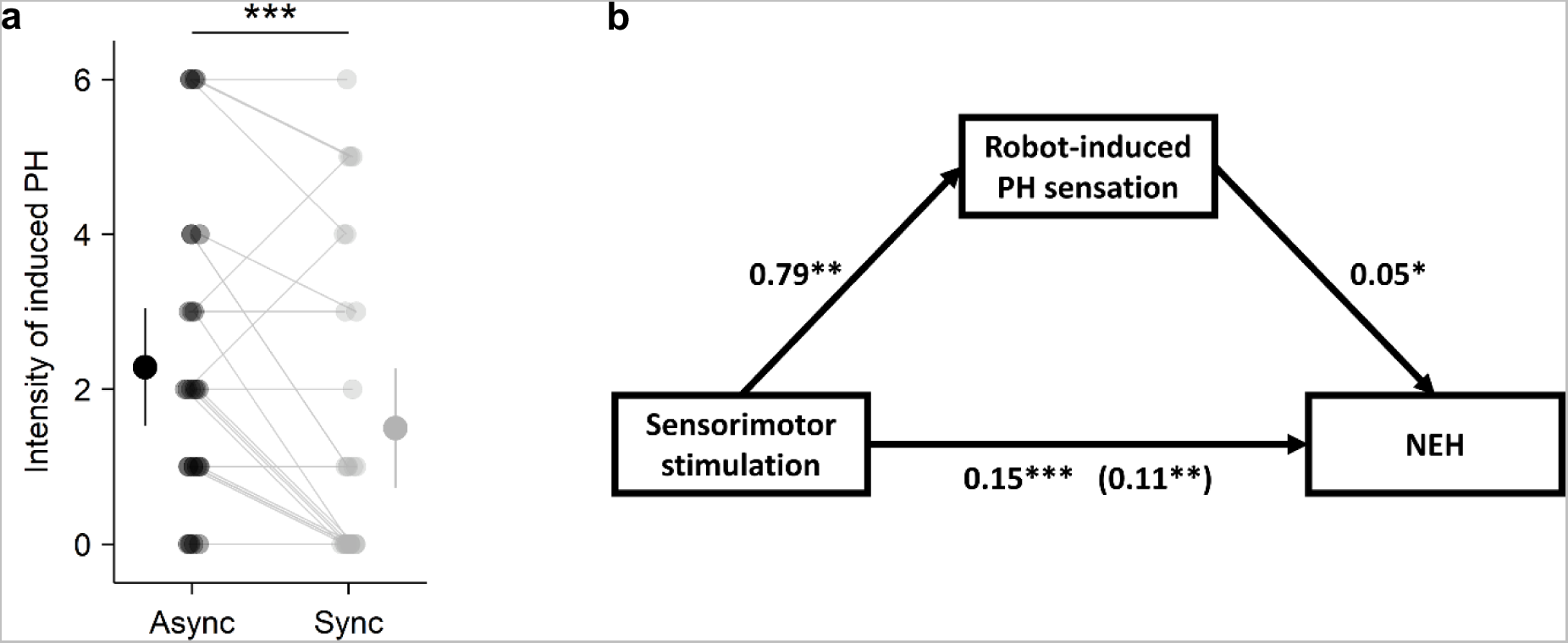
Robot-induced PH and its link to NEH (study 1). (a) robot-induced PH assessment ratings (asynchronous versus synchronous stimulation). Each linked pair of dots indicates the individual mean rating of robot-induced PH (asynchronous condition (dark grey) and synchronous condition (light grey)). The dots with the bar on the left and right sides indicate the mixed-effects linear regression between asynchronous (dark grey) and synchronous (light gray) sensorimotor stimulation. Error bar represents 95% confidence interval. (b) Results of mediation analysis. Standardized regression coefficients for the relationship between robotic sensorimotor stimulation, NEH, robot-induced PH sensation. *P ≤ 0.05; **P ≤ 0.01; ***P ≤ 0.001.

#### Overestimation of visual digital humans correlates with PH magnitude

To corroborate that NEH is an implicit marker for PH, we conducted additional correlation analysis between PH ratings and the magnitude of overestimation in NEH. This analysis revealed that the effect of sensorimotor stimulation on the NEH was partially mediated via PH ratings (Figure 4b, Note S2). This result indicates that the stronger our participants experienced PH during the asynchronous (versus synchronous) sensorimotor condition, the higher was their overestimation of digital humans shown in the virtual room (NEH). This finding was absent between PH ratings and NEO: the same analysis for NEO task during sensorimotor stimulation did not reveal any association between PH and NEO overestimation (Note S3), further confirming the selectivity of NEH overestimation as a marker for the proneness to experience a PH.

#### Summary of study 1

The main finding from study 1 is that we characterize NEH overestimation as a hallucination marker, showing that asynchronous sensorimotor stimulation is associated with an increase in NEH. Critically, this effect was observed (1) when comparing the PH-inducing asynchronous condition with the synchronous control condition, (2) was absent in the NEO task, and (3) cannot be explained by differences in task difficulty or alertness. Observing that (4) the magnitude of the NEH bias, but not the NEO bias, correlates with PH ratings, further links NEH overestimation with PH. Accordingly, we argue that a robotically induced mental state (i.e., the induction of a hallucinatory invisible percept) systematically modulates performance in a visual task (NEH) by increasing the magnitude of the number of seen humans. NEH is a robust marker for a specific and clinically relevant hallucination, PH, and is elicited in a controlled laboratory setting without relying on verbal ratings. NEH can be repeated as many times as needed, across control conditions, with full control over the presented stimuli. Integrated into fully automatized and a virtual training and test scenario, NEH thereby overcomes many limitations of previous hallucination research (Bernasconi, Blondiaux, et al. 2022; Rogers, Keogh, and Pearson 2021).

### Online numerosity estimations of visual digital humans reveals PH in Parkinson’s disease (study 2)

#### NEH in PD patients

These results show that NEH overestimation indexes experimentally induced PH in healthy participants. Would this quantitative, digital, and implicit maker for PH also extend to neurological patients, who experience PH as part of their disease? Would performance of PD patients with PH be characterized by an increase in NEH, compared to those without PH, even without any robotic stimulation? Recent work adapting the robotic hallucination-induction paradigm to PD indicated that PD patients experiencing symptomatic PH (PD-PH) had a six-fold higher sensitivity to the robotic sensorimotor procedure as compared to PD patients who never had PH (PD-nPH) (Bernasconi et al. 2021). In combination with the results from study 1, these clinical data suggest that (1) PD-PH patients may have a bias in NEH, that (2) this bias should exist without being exposed to robotic stimulation, and that (3) such a NEH bias should be larger than the one in PD patients without such hallucinations (PD-nH).

In study 2, we aimed to test NEH online and at home in a large cohort of PD patients in order to investigate whether NEH reveals the occurrence of PH in patients with spontaneous hallucinations and PD. This would be an important achievement, as it would allow to test patients directly at home, again without the biases associated with explicit verbal questionnaire evaluations or interviews (Rogers, Keogh, and Pearson 2021; Nisbett and Wilson 1977). Easy to use and unobtrusive online methods have gained momentum, by showing that large patient groups can be sampled, demonstrating the feasibility and validity of online methods and opening new perspectives towards better diagnostics and monitoring of diseases such as PD (Hillel et al. 2019; Brodie et al. 2016; Burq et al. 2022; Heijmans et al. 2019; Weil et al. 2018). Digital online NEH testing also facilitates reaching people living far away from medical centers (Galsky et al. 2015), in low income countries, without specific equipment (i.e., robotics, VR) and trained staff to perform hallucination testing.

We thus designed an online NEH task by adapting the method used in study 1 (immersive 3D VR paradigm integrated with robotics) to a web-based 2D task for NEH and for NEO (control task) without either VR or robotic stimulation. PD patients performed this web-based digital task at home, by themselves on their own personal computer or tablet. In addition to NEH and NEO data, we also acquired online questionnaire data about a range of demographical characteristics about the participants and about the occurrence of their hallucinations in daily life.

#### Demographical and clinical data

A total of 170 PD patients (Note S5) participated in our online experiment (https://go.epfl.ch/alpsn5, testing was carried out from August 2021 to June 2022) (Figure S13). From these, we included a total of 118 PD patients in the current analysis: 63 PD patients with PH (PD-PH) and 55 PD patients without any hallucinations (PD-nH) (see Methods). Analysis of demographic data did not show any significant differences in gender, age, disease duration, nor medication (Levodopa equivalent daily dose) between the two patient groups (i.e., PD-PH vs. PD-nH; Table 1).

**Table 1.** Clinical variables (study 2). Clinical variables of PD-PH and PD-nH included in the numerosity estimation task analysis. Of the 170 patients with PD, 118 patients of interest to answer our research question (PD-PH (n=63) and PD-nH (n=55)) were kept for the analysis of the NEH and NEO (see methods for further detail). The table shows the mean and standard deviation for several clinical and demographic variables.

|  | PD-nH (N=55) | PD-PH (N=63) | p-value |
| --- | --- | --- | --- |
| Gender | 27 (M) | 29 (M) | 0.7 (χ<sup>2</sup>) |
| Age | 66.11 ± 8.14 | 64.59 ± 7.57 | 0.30 |
| PD duration (year) | 6.18 ± 5.26 | 6.42 ± 4.95 | 0.80 |
| LEDD (mg) * | 276.56 ± 317.20 (N=53) | 318.49 ± 315.11 (N=61) | 0.48 |

#### NE for 2D human stimuli in a home-based online setting using participants’ personal computer

To test NE for different numerosities of humans and to evaluate whether classical NE effects can also be observed for human stimuli on a 2D screen (as observed previously for visual dots) (Arrighi, Togoli, and Burr 2014; Cicchini, Anobile, and Burr 2014; Fornaciai and Park 2020; Jevons 1871; Kaufman et al. 1949; Poulton 1979; Minturn and Reese 1951; Krueger 1984) and for 3D digital humans (i.e., study 1), we converted our 3D VR stimuli, used in the NEH and NEO tasks of study 1, into 2D stimuli that we displayed on participants’ computer screen or tablet at home. We presented a varying number of digital humans (Figure S1) that were equally positioned across the depicted room, characterized by the same configurations as used for the VR stimuli of study 1. Following a screen calibration procedure to control the size of the displayed stimuli, each participant performed the online NE task. Based on previous NE work, participants were asked to indicate the number of humans (online NEH task) they perceived in the room, as fast and as precise as possible. For the control condition (NEO) the digital humans were replaced with objects (boxes; online NEO task, Figure S1), positioned in the same way as the digital humans. In the control NEO task, participants were asked to indicate the number of objects they perceived in the room.

First, as predicted based on previous literature (Jevons 1871; Kaufman et al. 1949; Minturn and Reese 1951) and on the data of study 1, online data show that NE is modulated by the number of stimuli (humans or objects) presented in the virtual room (i.e., presented numerosity; F(3, 9084)=2055; p<0.001; main effect; table S5). In particular, we observed that NE increases with the number of presented numerosities, independently of the type of stimulus (Table S5; Table S9; Table S10). Second, additional post-hoc analysis showed that participants mean NE is significantly higher than the presented numerosity in the range of presented numerosities (5 to 8) (Figure 6; Table S5). This is a typical behavior observed with dots stimuli for numerosity just above the subitizing range (Fornaciai and Park 2020; Jevons 1871; Minturn and Reese 1951). These data extend two well-known known effects from NE studies that have been carried out in research laboratories in young healthy participants (numerosity main effect, overestimation just above the subitizing range) for the first time to elderly PD patients, who carried out the task on their personal computer or tablet at home.

**Figure 5.**
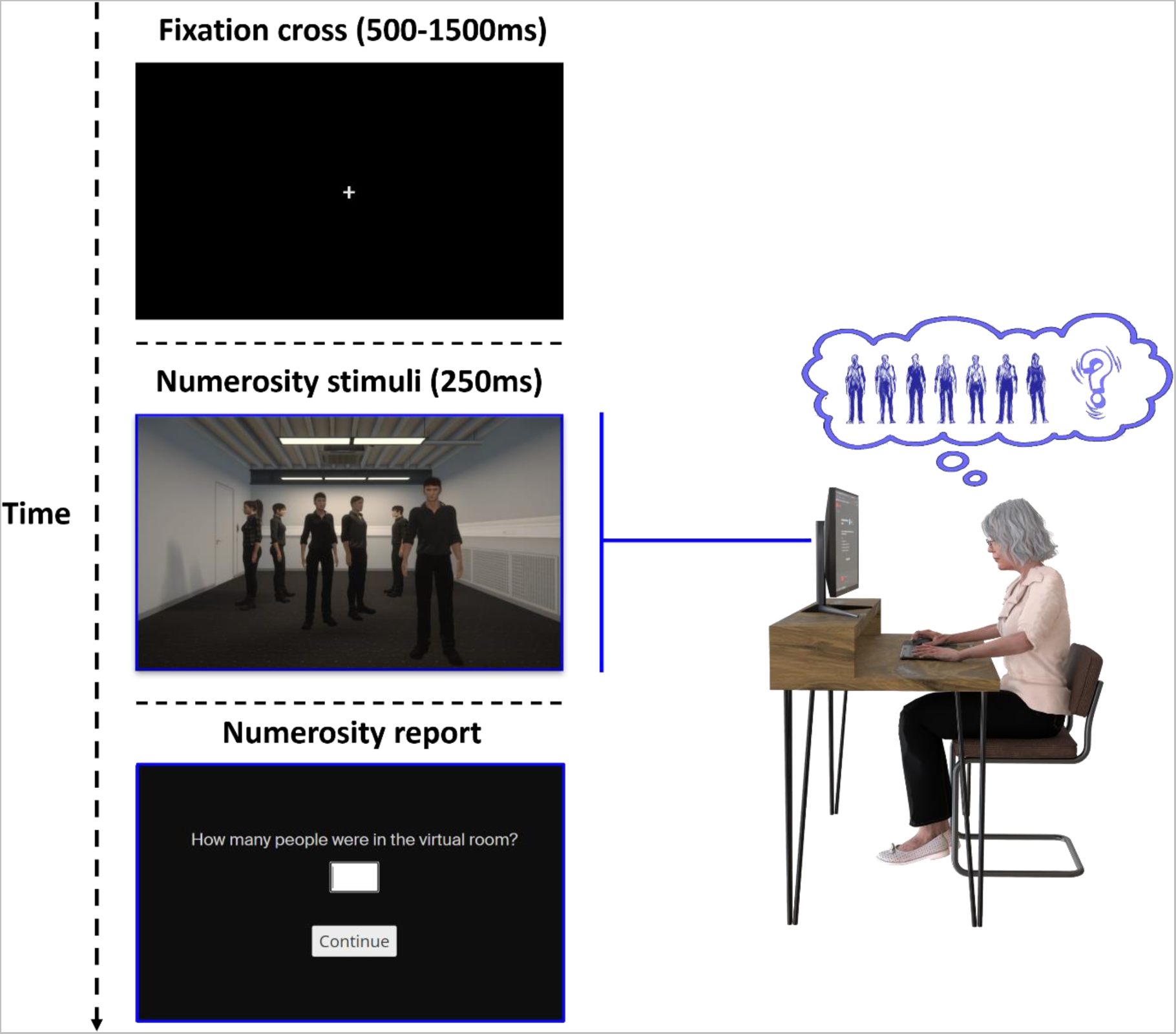
Online NEH task (study 2). A single online NEH trial is shown. that started with the appearance of a fixation cross (500-1500ms). After that a scene containing different number of people (range 5-8) was shown for 250ms in a visual scene on their screen and PD patients were asked to estimate the number of people that they saw. PD patients performed this web-based digital task at home, on their personal computer or tablet.

**Figure 6.**
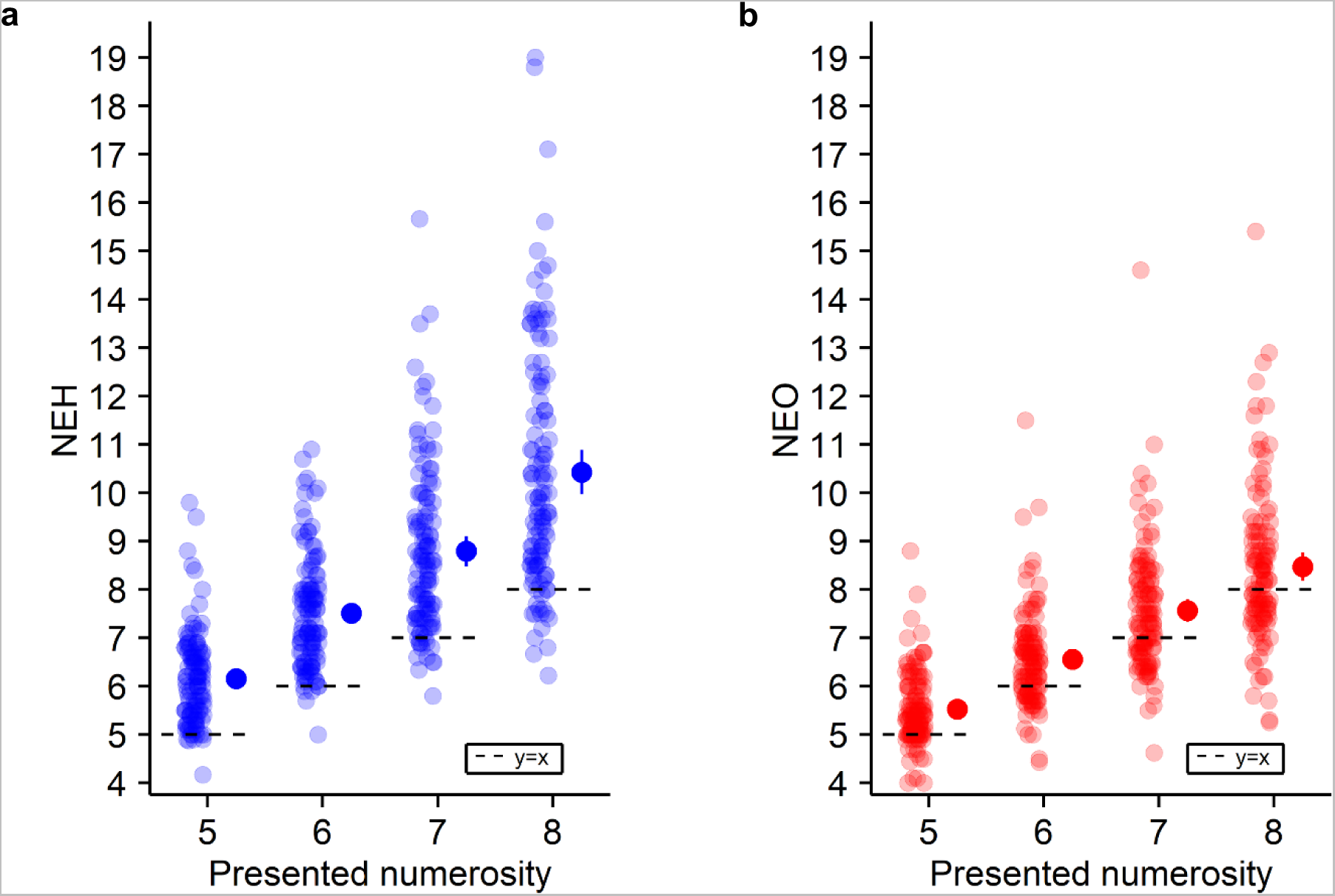
Numerosity estimation (study 2). General numerosity estimation performance for each tested numerosity in the (a) NEH and (b) NEO task (study 2). Each dot indicates the individual NEH mean (over the trials) estimate at the corresponding tested numerosity. The dots with the bar on the right sides indicate the in-between subject mean at each presented numerosity. Note the general overestimation bias in NEH and NEO. Error bar represents 95% confidence interval.

#### PH in patients with PD is associated with higher overestimation in NEH

Our main research questions were whether the occurrence of PH in PD (PD-PH group) is associated with changes in online NEH and, in particular, if PH is associated with overestimation in online NEH, as observed in healthy participants during robot-induced PH (study 1). Additionally, based on the findings of study 1, we predicted that such an effect would be absent in the online control NEO task (Figure S1). In agreement with our hypothesis, we observed an overestimation in the PD-PH group as compared to the PD-nH group. Moreover, overestimation was only present for the online NEH, but absent for the online NEO task. Indeed, our results show that online NE is significantly (F(1, 9085)=53.81; p<0.001; interaction; Table S9) modulated by the occurrence of PH (PD-PH vs PD-nH) and by the type of stimulus (online NEH vs. online NEO). Critically, and in agreement with our hypothesis, post-hoc comparisons showed that the occurrence of PH significantly modulates online NEH (t(122)=−3.16; p=0.002; Table S9) (Figure 7a; Figure 7b), with PD patients who reported PH showing stronger online NEH overestimation. Moreover, this effect was only present when estimating the number of humans in the virtual room and was absent for online NE for control objects: further post-hoc analyses did not reveal any difference in online NEO between PD groups (PD-PH vs PD-nH) (statistical t(122)=−0.59; p=0.42; Table S9) (Figure 7c and Figure S10). These data show that PD patients with PH show an NEH overestimation bias that can be measured online at the patient’s home. We also corroborate the specificity of the overestimation bias for human stimuli, because the effect was absent in the online NEO task, showing that NEH is an implicit digital online marker for PH in PD patients.

Additional analysis revealed no statistical differences in response times between the two PD groups (i.e., PD group; F(1,116) = 0.58; p=0.45; no main effect nor any interaction; Table S11) (Figure S8; Figure S9; Figure S11; Figure S12), showing that task difficulty (online NEH and online NEO) did not differ between both PD groups (PD-nH and PD-PH). These results consolidate our previous findings and show that the overestimation observed in PD-PH versus PD-nH and in the online NEH versus online NEO cannot be explained by differences in task difficulty between groups.

#### Summary of study 2

We successfully adapted our previously developed method of study 1 (immersive VR NE task on humans and control objects; robotic sensorimotor stimulation) into a digital procedure that is fully online, does not require any robotic stimulation, and engaged PD patients at their homes performing the task on their personal computer or tablet (the dropout rate during the NE tasks was only 7%; reasons for participant drop-out are indicated in Supplementary Note 8). Applying this new online procedure, we, critically, show that the online performance of PD-PH patients is characterized by a stronger online NEH overestimation bias (as compared to PD patients without hallucinations; PD-nH), thereby linking clinical PH to online NEH overestimation. This effect was absent in the online NEO task and cannot be explained by differences in task difficulty between groups, nor does it depend on clinical covariates such as age, gender, disease duration, or first affected side, corroborating data from study 1, extending them to a large clinical cohort of PD patients, and underlining NEH overestimation as a digital marker of PH in PD.

## Discussion

We developed a new paradigm (Numerosity Estimation of Humans – NEH) to implicitly assess a clinically-relevant hallucination: PH. In a first study in healthy participants, we tested our paradigm by merging immersive VR with a robotic platform able to experimentally induce PH in a controlled manner and allowing for the real-time investigation of PH (Bernasconi, Blondiaux, et al. 2022; Bernasconi et al. 2021; Blanke et al. 2014). Our results show that robot-induced PH are associated with a selective overestimation for digital humans (NEH), but not for control objects (NEO). These results confirm our hypothesis that NEH is a robust digital marker for the specific hallucinatory mental state of PH. Because hallucinations are frequent and clinically relevant phenomena in PD, in a second study, we adapted our NE task to a web-based digital test and investigated 170 PD patients remotely at their homes. Translating our procedure to an online assessment, we validate NEH overestimation as a digital marker for PH occurring as a symptom in PD. Using several controls, we rule out the possibility that these NEH effects are confounded by clinical variables or task demands.

In study 1, we characterize NEH overestimation as a hallucination marker by showing that the PH-inducing robotic sensorimotor condition is associated with a higher NEH (Figure 3a, Figure 3b), linking a robot-induced mental state with NEH overestimation. Importantly, this effect is absent for NEO (Figure 3c) and cannot be explained by differences in task difficulty or attentional processes. We further report a correlation between the magnitude of the NEH overestimation and PH ratings (Figure 4b), corroborating the link between NEH overestimation with PH. These results suggest that the robot-induced PH state systematically modulates the performance in NEH by increasing the magnitude of the number of people perceived. Compared to previous implicit measures used to assess PH (e.g. drift in self-location (Blanke et al. 2014), numerosity of actual people close by (Blanke et al. 2014)), the present NEH has several advantages. The presented human stimuli are fully controlled, are tested for different and larger numerosities, and based on many repeated trials. Critically, the present procedure encompasses a carefully matched control condition with non-human digital objects (NEO), and we found no asynchrony-dependent overestimation when our healthy participants judged the number of non-human stimuli (NEO), controlling for task demands. Finally, NEH can be repeated numerous time for all experimental stimuli and conditions and is fully orthogonal to the robotic sensorimotor manipulation, overcoming limitations of previous hallucination research (Bernasconi, Blondiaux, et al. 2022; Rogers, Keogh, and Pearson 2021) and indexing a specific and clinically relevant hallucination in PD: PH.

In study 2, we successfully translated NEH measurements to an online assessment of hallucinations performed by elderly PD patients, who carried out the task on their personal computer or tablet at home. Data show that PD patients with disease-related PH in their daily life have a stronger online NEH bias and overestimation, as compared to PD patients without any hallucinations (Figure 7a, Figure 7b). This effect is absent for the online NEO assessment (Figure 7c), cannot be explained by between-group differences in task difficulty, and does not depend on any of the acquired clinical covariates (i.e., age, gender, disease duration, first affected side). The online NEH bias in PD-PH (study 2) thus extends the NEH bias, as induced by asynchronous robotic stimulation in healthy individuals (study 1), to patients with PD. We argue that online NEH is a robust and quantitative digital marker for PH in PD and, potentially, for other hallucinations and for cognitive decline, because PH share brain alterations with visual hallucinations (Pagonabarraga et al. 2014; Helena Bejr-kasem et al. 2019) and have been linked to cognitive impairment (Bernasconi et al. 2021) and more rapid cognitive decline (Bernasconi, Pagonabarraga, et al. 2022; H. Bejr-kasem et al. 2021). Compared to current standard methods used to assess hallucinations in the clinic (based on verbal self-reports and interviews of patients and interpretations by clinicians), which are associated with well-known biases, our implicit online NEH-based assessment overcomes several methodological limitations such as participant and experiment biases and underreporting due of fear of stigmatization (Rogers, Keogh, and Pearson 2021; Nisbett and Wilson 1977; Bernasconi, Blondiaux, et al. 2022). Thus, the present online NEH measure constitutes a new promising digital marker for the quantitative assessment and monitoring of a more severe form of PD, associated with hallucinations as well as cognitive decline (Masanneck et al. 2023; Stroud, Onnela, and Manji 2019). The present findings add to the recent upsurge of the impact of digital health technologies in PD (Mahadevan et al. 2020; Burq et al. 2022). These studies collected digital measures focusing on motor symptoms and their fluctuations (i.e., ~70% of studies on motor function vs. ~10% on cognitive function) (Masanneck et al. 2023), whereas the present NEH findings constitute, to the best of our knowledge, the first online digital health marker for hallucinations in PD. Easy to use and unobtrusive online methods allow the investigation of much larger patient groups and facilitate longitudinal sampling, enabling better diagnostics and monitoring (Hillel et al. 2019; Brodie et al. 2016; Burq et al. 2022; Heijmans et al. 2019). Clinic-based versus home-based evaluations often provide only a single snapshot of patient performance and this may not properly reflect a patient’s performance in daily life at home (Takayanagi et al. 2019; Hillel et al. 2019; Brodie et al. 2016). The present digital online NEH testing has the additional advantage of reaching people living far away from medical centers, in low income countries, without requiring any specific equipment or trained staff (Galsky et al. 2015; Burq et al. 2022).

**Figure 7.**
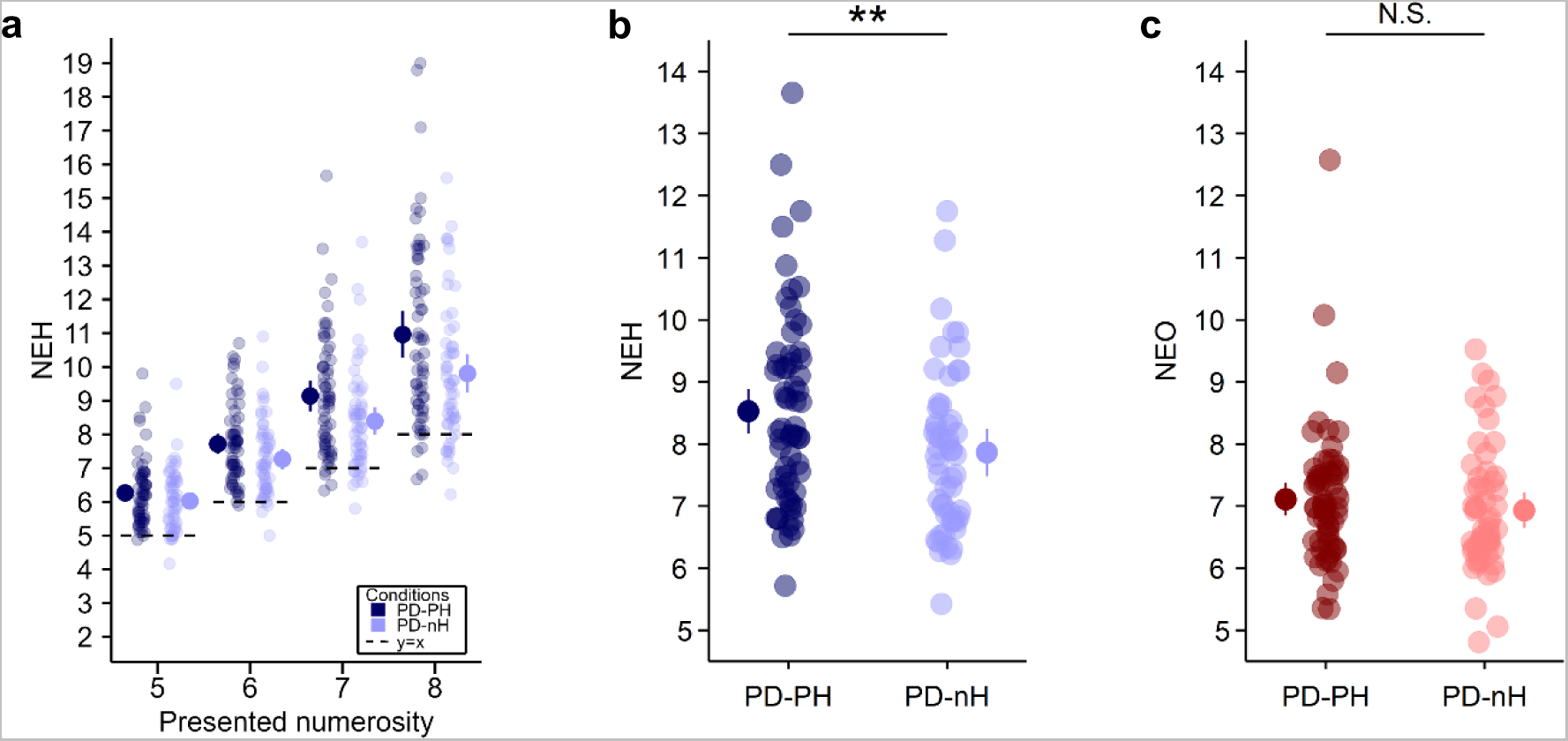
NEH and NEO for both PD patient groups (PD-PH and PD-nH) (study 2). (a) Performance is shown in PD patients for each tested numerosity in the NEH task for PD-PH and PD-nH separately. Each dot indicates the individual NEH mean estimate for the tested numerosity (PD-PH (dark blue) and PD-nH (light blue)). The dots with the bar on the left and right sides indicate the mixed-effects linear regression between PD-PH (dark blue) and PD-nH (light blue) at each presented numerosity. (b) NEH in PD patients (PD-PH vs PD-nH). Each dot indicates the individual NEH mean estimate (PD-PH (dark blue) and PD-nH (light blue)). The dots with the bar on the left and right sides indicate the mixed-effects linear regression between PD-PH (dark blue) and PD-nH (light blue). (c) NEO in PD patients (PD-PH vs PD-nH). Each dot indicates the individual NEO mean estimate (PD-PH (dark red) and PD-nH (light red)). The dots with the bar on the left and right sides indicate the mixed-effects linear regression between PD-PH (dark red) and PD-nH (light red). Error bar represents 95% confidence interval. **P ≤ 0.01.

Concerning virtual, augmented and mixed reality in medicine, immersive VR has recently emerged as a prominent tool for supportive treatment in mental health disorders (Maples-Keller et al. 2017) and rehabilitation (Georgiev et al. 2021), with recognized methodologies for validating its use for clinical interventions (Birckhead et al. 2019). The use of VR as a diagnostic tool, however, remains relatively underexplored (Moon and Han 2022), despite promising studies in mental health and cognitive function (Freeman et al. 2017; Riva and Serino 2020). Moreover, the present procedure in experiment 1 is based on the integration of immersive VR and robotics, allowing us to go beyond previous findings (Bernasconi et al. 2021; Blanke et al. 2014; Orepic et al. 2021; Serino et al. 2021; Salomon et al. 2020), by describing classical NE effects for complex scenes with 3D digital humans and 3D objects (Fornaciai and Park 2020; Jevons 1871; Minturn and Reese 1951) and by revealing their dependence on PH. We also show that immersion in VR allows the reproduction of the same exposure and experimental conditions, for all participants, as immersive VR allows to provide the task instructions in a pre-recorded yet interactive manner, thereby automatizing the experimental procedure (i.e., the 3D digital experimenter provides the experimental instructions in VR, physical interactions with the real experimenter are minimized) and limiting potential experimenter biases (Horing et al. 2016; Moon and Han 2022; Rosenthal 1976; Doyen et al. 2012). Moreover, because the exact same VR program can be executed by other researchers and in other settings, the experiment can be reproduced with minimal differences, thus enhancing the comparability between studies by different research groups and improving double-blind experimental designs and multi-center studies (Horing et al. 2016).

There are several limitations of our study. First, in study 1, although the overestimation bias was found in the NEH, but not the NEO task, we only compared the effects of 0ms (synchronous) and 500ms delay (asynchronous) sensorimotor conflicts on PH. As robot-induced PH has been shown to depend on the degree of sensorimotor conflict (Bernasconi et al. 2021), future studies should test multiple delays of sensorimotor conflict (Bernasconi et al. 2021), further defining the link between the intensity of robot-induced PH and the overestimation of digital humans. Second, in study 1, participants are immersed in a virtual environment where a virtual character speaks to them (the virtual experimenter) and where several others enter and leave the room where the participant feels present. Our manipulation leverages on the subjective experience of copresence (togetherness with others in the virtual world (Slater et al. 2022)) that occurs in such VR simulations. Although copresence is often observed in VR with similar settings and rendering quality, our study could benefit from a direct assessment of copresence. Third, the data of study 2 were acquired through an anonymized home-based online measurement, with limited information regarding the patients’ symptoms and other clinical variables. Future clinical work should explore the novel digital PH marker jointly with detailed neurological, neuropsychological and other clinical evaluations. Fourth, we carried out a cross-sectional study. Future work may also perform longitudinal assessments investigating for example the stability of the NEH marker over time, whether it reflects hallucination frequency, and how it relates to other hallucinations.

In conclusion, the present data demonstrate that VR NEH and online NEH are new digital markers for PH, in healthy individuals as well as in patients with PD. By merging robotics, VR technology, and NE, we report that experimentally induced PH in healthy participants results in NEH overestimation (but not in the control NEO task), revealing that NEH is a quantitative and robust digital marker for an experimentally induced hallucinatory mental state. By translating this VR-based marker to a home-based online assessment, in patients with PD, we show that NEH overestimation is a digital marker for disease-related PH that PD patients experience in their daily lives. Using a combination of controls, we ruled out the possibility that these effects stem from clinical variables or task demand. The present digital marker advances previous hallucination evaluations, as based on verbal interviews, by proposing a robust and quantitative assessment of these subjective mental states that are highly prevalent in many neurological and psychiatric diseases (Bernasconi, Pagonabarraga, et al. 2022). Our online home-based procedure strongly improves accessibility that often limits the impact of new laboratory-based markers and allows to reach people living far away from medical centers without requiring any specific equipment or trained staff (Galsky et al. 2015; Burq et al. 2022). As there is growing evidence that specific hallucinations, such as PH, are an early marker for cognitive decline and dementia in PD (Bernasconi, Pagonabarraga, et al. 2022; Bernasconi et al. 2021; H. Bejr-kasem et al. 2021; Lenka et al. 2019), NEH may not only detect proneness for psychosis, but also for later cognitive decline, requiring further longitudinal work.

## Methods

### Experiencing an invisible presence increases visual overestimation bias for numerosity estimations of visual digital humans (study 1)

#### Preregistration

The study preregistration is available at https://doi.org/10.17605/OSF.IO/YR3CP (Albert et al. 2021)

#### Study population

Twenty-height healthy participants (18 women, 10 men; age ranging from 18 to 33 years, mean ± SD age = 24 ± 3.42 years) took part in this experiment. They were all right-handed according to the Edinburgh Handedness Inventory(Oldfield 1971) (score ranging from 50 to 100, mean ± SD score = 87.5 ± 18). All the participants signed a written informed consent before participating in the experiment and were rewarded for their time with monetary compensation (CHF20/hour). None of the participants had current nor history of neurological, psychiatric and substance abuse disorders. Participants were also screened for a good stereoscopic vision threw a stereoscopic acuity test described in Albert et al. (Gauthier et al. 2021). All participants included in the study were naive to the purpose of the experiment. The experimental procedures (under protocol reference n° 2015-00092) were approved by the Cantonal Ethics Committee of Geneva (Commission Cantonale d’Ethique de la Recherche sur l’Être Humain - CCER).

#### Apparatus and materials

##### Robotic system

The robotic system is composed of a commercial haptic interface (Phantom Omni, SensAble Technologies), coupled with a custom three degree-of-freedom robot in the back (Hara et al. 2011). Participants are sitting on a chair and controlling the front robot situated on a table directly in front of them with their right index finger. The back robot is located behind their back and reproduces the movements initiated with the front robot with virtually no delay in the synchronous condition, and with 500 msec delay in the asynchronous condition, which has been shown to induce PH. This creates different degrees of sensorimotor conflict between the right-hand movement and the somatosensory feedback on the back (Blanke et al. 2014).

##### Virtual Reality system

The virtual reality system consists in a commercial Head Mounted Display system (Oculus Rift CV1 coupled with a single Oculus Rift Sensor).

##### Auditory system

The auditory system consists of a commercial Auditory headset (Arctis Wireless Pro).

##### Graphic system

The graphic system consists in a laptop computer running Windows 10 and equipped with an Intel i7 6700HQ processor, 16Gb Random-Access Memory, and a Nvidia GeForce GTX 1060 graphic card, ensuring a constant 90Hz display rate of the experiment in the Virtual reality system. The Virtual Reality system is plugged into the graphic system. The experiment running on the graphic system is implemented in Unity 2019.3.13f1 using C#.

##### Experiment control and progress tracking system

The experiment is controlled, and progress tracked in real time through a mobile-compatible application, implemented in Unity 2019.3.13f1 using C#. The device used consists in a smartphone running Android (Samsung Galaxy S8+), connected to the graphic system over Wi-Fi.

#### Digital experimenter

The digital experimenter was created with Character Creator and the Headshot plugin. The model of the digital experimenter is available as supplementary material (see Code availability section). All the instructions are given by the digital experimenter (Video S3; Video S4; Video S5), whose lip movements are automatically synchronized with the prerecorded audio using speech-driven lip-sync (Llorach et al. 2016).

#### Numerosity stimuli

Visual stimuli were generated on Unity 3D (version 2019.3.13f1). The virtual environment was modeled in 3DS max and consisted of a realistic representation of our experimental room. The model of the virtual environment is available as supplementary material (see Code availability section). The virtual scene exposure was set to high in order to have a low global brightness, to encourage hallucinations induction which have been reported to often be triggered in low levels of illumination in patients’ daily life (Chaudhury 2010; Lana-Peixoto 2014; Barnes and David 2001). Stimuli consisted of a 3D scene of our virtual experiment room with digital humans (or control objects) inside. Digital humans and control objects were placed in the virtual scene in front of the camera viewpoint, facing the viewpoint in a range from – 90° to 90°, and in a way that they do not overlap completely from the viewpoint. Digital humans and control objects were placed between 1.75m and 5.25m in depth from viewpoint, and between −1.5m and 1.5m from right to left. Array of digital humans and control objects occupied a maximum virtual camera visual angle of 60° horizontally. This ensures that digital humans and control objects were located within the participants’ peripheral field of view, which is the “vision produced by light falling on areas of the retina outside the macula” (Morris, Press, and Morris 1992), and that they were displayed inside and close to the limit of 30° retinal eccentricity, which is the limit from which the visual acuity decreases more strongly (Anstis 1974). The digital humans and control objects arrays were ranging from 5 to 8 and generated on the fly based on seeded configurations. The lower bound was decided as the lower bound of the estimation range of our type of stimuli (Note S1).

#### Robot-induced subjective experiences questionnaire

Participants performed the robot-induced PH paradigm for 2 minutes, one time in synchronous condition and one time in asynchronous condition, randomized order across participants. After each condition, a lab-tailored questionnaire was used to measure the presence hallucination (“I felt as if someone was standing close to me (next to me or behind me)”), along with other subjective experiences as self-touch (“I felt as if I was touching my back myself”), passivity experience (“I felt as if someone else’s was touching my back”), and loss of agency (“I felt as if I was not controlling my movements or actions”). Three control questions were also asked (“I felt as if someone was standing in front of me”, “I felt as if I had two body”, “I felt anxious/stressed”). These 7 questions measuring PH and other illusions were adapted from Blanke et al. (Blanke et al. 2014). At the beginning of the robot-induced PH paradigm, participants were put in a dark environment, and following an acoustic cue (beep, 1000Hz, 250msec) they were asked to close their eyes and start performing the poking movements with the front-robot. During the whole trial duration white noise was presented through headphones to isolate participants from the robotic noise. After 120sec a second acoustic cue (double beep, 1000Hz, 250msec each beep separated by 250msec) indicated the end of the trial, and participants were asked to answer questions displayed in the Head Mounted Display. The questions were displayed in a randomized order across conditions (synchronous and asynchronous) and participants. Participants were asked to indicate on a 7-point Likert scale how strongly they felt the sensation described by each item (from 0 = not at all, to 6 = very strong). The yaw orientation of the HMD was used to target a value on a slider. Once the desired value was targeted, participants were instructed to say verbally either “ok” or “validate” to validate their current answer. Validation was performed by voice recognition (Windows Speech KeywordRecognizer). An overview of the robot-induced subjective experiences questionnaire protocol is shown in Extended Data Figure 1. A complete overview of the virtual instructions is available in Video S3.

#### Numerosity estimation

The numerosity estimation task was divided into two parts, one presenting arrays of digital humans (NEH) in the virtual environment and the other one presenting arrays of control objects (NEO) in the virtual environment, randomized order across participants. Each part contained four blocks of respectively twelve, eight, eight and twelve trials. Blocks of twelve (respectively height) trials contained three (respectively two) times each numerosity. Each block started with a 60sec habituation phase, where several digital humans moved and discussed in the virtual environment in the digital human part (around 10 in total per habituation phase) (Video S6; Video S7; Video S8; Video S9). Some of them (between 2 and 3 depending on the habituation phase) were discussing together. There was a total of four different habituation phases, presented in the same order across participants. In the control objects condition, control objects replacing, and matching position and rotation of digital humans appeared and disappeared along time in the virtual environment (Video S10; Video S11; Video S12; Video S13). At the end of a habituation phase, trials started. At the beginning of a trial, participants were put in a dark environment, and following an acoustic cue (beep, 1000Hz, 250msec) they were asked to close their eyes and start performing the poking movements with the front-robot. The back robot sensorimotor stimulation was either synchronous (0ms delay) or asynchronous (500ms delay), alternating across blocks, starting condition equally balanced across participants. During the whole trial duration white noise was presented through headphones to isolate the participant from the robotic noise. After 30sec a second acoustic cue (double beep, 1000Hz, 250msec each beep separated by 250msec) indicated the end of the trial. Participants then had to open their eyes and keep their gaze on a central fixation cross, which was briefly presented (range between 500msec and 1500msec) before the presentation of the visual stimulus. The visual stimulus, consisting in an array of digital humans or control objects (ranging from 5 to 8) in the virtual environment, was then presented in front of the participant for a duration of 200msec. Participants then had to report the number of digital humans (“How many people are in the room?”) or control objects (“How many objects are in the room?”) they estimated to be in the virtual environment on a scale ranging from 0 to 20. As in task 1, the yaw orientation of the HMD was used to target a value, and voice recognition (Windows Speech KeywordRecognizer) was used to validate the answer. There was a total of 10 different stimuli configurations possible per numerosity (ranging from 5 to 8), where stimuli type (human or control objects) had a specific configuration (in terms of stimuli position and orientation, which were matched between digital humans and control objects for each configuration). Half of the participants were associated with 5 stimuli configurations per numerosity, the other half of the participants were associated with the 5 remaining stimuli configurations per numerosity. An overview of the numerosity estimation protocol is shown in Extended Data Figure 1.

#### Data analysis

##### Robot-induced sensations questionnaire

Cumulative link mixed model (packages ordinal (Christensen 2019) and RVAideMemoire (Hervé 2022) in R (Team 2013)) were used to analyze questions of the robot-induced sensations questionnaire. Sensorimotor stimulation (synchronous or asynchronous) was set as fixed effect, and random intercept for each subject were assumed. The significance of fixed effects was estimated with likelihood Ratio Test comparing full model (with sensorimotor stimulation set as fixed effect) against a reduced model without the fixed effect in question. We concluded that the fixed effect (condition) was significant if the difference between the likelihood of the two models was significant.

##### Numerosity estimation

Linear mixed effects models (packages lme4 (Bates et al. 2015, 4) and lmerTest (Kuznetsova, Brockhoff, and Christensen 2017)) with robotic sensorimotor stimulation (synchronous or asynchronous), presented numerosity and type of stimuli (human and control objects) as fixed effect, random intercept for each subject was performed on the numerosity estimation data. The significance of fixed effects was estimated with likelihood Ratio Test. Post-hoc analysis was performed on the significant interactions and corresponded in pairwise comparisons using independent-samples t-tests, corrected with Holm’s sequential Bonferroni procedure.

The general estimation performance at each numerosity was assessed with one-sample t-tests against presented numerosity, with reported p-values not corrected for multiple comparisons. The difference of estimation between NEH and NEO was assessed with two-sample t-tests at each of the presented numerosity, with reported p-values not corrected for multiple comparisons.

Trials in which participants did not see the stimuli (answer 0 or 1) were excluded. This procedure resulted in the exclusion of no trials across conditions and participants.

### Online numerosity estimations of visual digital humans reveals PH in Parkinson’s disease (study 2)

In this online web-based experiment, participants first filled in some socio-demographic information, answered a questionnaire on alteration of perception corresponding to the frequency of hallucination occurrence in daily life, followed by a screen calibration procedure, and the numerosity task. The experiment was available in French and English. The experiment could be performed on a computer or on a tablet.

#### Study population

One hundred and seventy patients with PD (93 women, 77 men; age ranging from 42 to 79 years, mean ± SD age = 65.4 ± 7.83 years; PD duration ranging from 1 month to 25.6 years, mean ± SD PD duration = 6.44 ± 5.19 years) took part in this study. Of the 170 patients with PD, 118 patients of interest to answer our research question were kept for the analysis of the NEH and NEO task in the current paper (see Data analyses section below for details).

#### Socio-demographic information

At the beginning of the experiment, participants were asked to indicate their gender, age, country, time, visual disturbances, and whether they had been diagnosed with Parkinson disease. In case of positive answer to this last question, participants were also asked the date of diagnosis (year and month), the side of the body where symptoms appeared first (“left”, “right”, “both”, “I don’t know”), the current medication along with the daily dosage, and the time of the last medication intake for Parkinson’s disease (Levodopa).

#### Questionnaire on alteration of perception

This questionnaire is a self-assessment questionnaire on whether the frequency of occurrence of specific hallucinations in daily life (passage hallucination, presence hallucination, visual illusion, or visual hallucination – see Table S6 for the list of questions). Participants answered on a 5-item Likert scale (0 – Never; 1 – Rarely (less than once a month); 2 – Occasionally (several times, but less than once a week); 3 – Frequently (several times a week, but less than once a day); 4 – Daily (almost every day, several times a day); see Table S8 for the report of these occurrences).

#### Screen calibration

Participants were invited to measure and report the length of a line displayed on their screen. This measure allowed to scale stimuli, so they were displayed the same physical size on each participant screen, independent of monitor screen and positions. Participants were instructed to stay one meter away from their screen on a computer and fifty centimeters away from their screen on a tablet. The detection of whether the participants were using a computer, or a tablet was automatic.

#### Numerosity task

The numerosity estimation task was divided into two parts, one NEH and one NEO, randomized order across participants. Each part contained 40 trials, 10 trials per numerosity (ranging from 5 to 8), randomized order across participants. There was a total of 10 different stimuli configuration possible per numerosity (ranging from 5 to 8), which were the same as in the behavioral experiment (Numerosity estimation as an implicit measure for robot-induced PH). The stimuli were 1280px width by 720px height pictures and measured onscreen approximately 35.2 × 19.8 cm on the computer version and 17.6 × 9.9 on the tablet version. The stimuli were displayed for 250ms(Cicchini, Anobile, and Burr 2014; Fornaciai and Park 2020; Birren and Botwinick 1955).

#### Apparatus and material

This online experiment was developed in house using javascript and the jsPsych library(de Leeuw 2015) (on client side: javascript, html, css; on server side: https server in node.js, nginx as a reverse proxy, running in two docker containers) and was hosted on an EPFL Server in a dedicated Virtual Machine (1xvCPU, 1GB RAM, 40GB HDD) in Demilitarized Zone. Data were saved and stored on EPFL’s servers.

#### Data analysis

##### Numerosity estimation

Of the 170 patients who participated in our online study, we included a total of 118 PD patients in the analysis of the numerosity estimation task: 63 PD patients with PH (PD-PH) and 55 PD patients without any hallucinations (PD-nH). The selection criteria are described below. Participants who had a very low refresh rate (less than 20Hertz) or resolution (less than 800px on both axis) were excluded from the analysis of the numerosity task. This resulted in the exclusion of respectively 4 and 4 participants. Participants who reported visual disturbances that could negatively impact the task were also excluded from the analysis of the numerosity task (mainly diplopia, the reports of visual disturbances of these participants can be found in Table S7). This resulted in the exclusion of 10 participants. Participants who reported hallucinations, but not PH (passage hallucinations, visual illusions or structured visual hallucinations, n = 39) were also excluded from the current analysis. In the selection procedure described above, some participants belong to different categories of rejection criteria.

Trials in which participants did not see the stimuli or made an evident mistake in their reporting (answer less or equal to 3 or answer superior to 50), along with trials in which participants took too much time to give an answer (response time superior to 15seconds) were excluded from the analysis of the numerosity task. This procedure resulted in the exclusion of 2.2% of the trials in the NEH task (104 trials over 4720) and 2.5% of the trials in the NEO task (120 trials over 4720).

Participants were then separated into two groups based on their response to the questionnaire on alteration of perception (Table S6). The first group (PD-nH group) contains PD patients not having experienced any kind of hallucination (response equal to 0 to all questions). The second group (PD-PH group) contains PD patients having experienced PH (response greater than or equal to 1 to the corresponding question). Demographic and clinical characteristics of the PD-nH and PD-PH group population are reported in Table 1.

Linear mixed effects models with PD group (PD-nH or PD-PH), presented numerosity and type of stimuli (digital humans and control objects) as fixed effect, random intercept for each subject was performed on the numerosity estimation data. The significance of fixed effects was estimated with likelihood Ratio Test. Post-hoc analysis was performed on the significant interactions and corresponded in pairwise comparisons using independent-samples t-tests, corrected with Holm’s sequential Bonferroni procedure.

The general estimation performance at each numerosity was assessed with one-sample t-tests against presented numerosity, with reported p-values not corrected for multiple comparisons.

##### Clinical variables

In Table 1, gender independence was compared between groups (PD-nH vs PD-PH) with a chi-square test. Age, PD duration and Levodopa equivalent daily dose (LEDD) were compared between groups (PD-nH vs PD-PH) with two-sample t-tests. LEDD was calculated as a sum of the conversion of each parkinsonian medication to Levodopa Equivalent Dose (Nyholm and Jost 2021; Schade, Mollenhauer, and Trenkwalder 2020; Tomlinson et al. 2010). Four PD patients did not correctly report their medication (two PD-nH and two PD-PH); thus, they were excluded from the calculations and statistical test of LEDD.

## Data availability

The main data supporting the results in this study are available within the paper and its supplementary Information. The source data along with the analysis scripts have been deposited in https://gitlab.epfl.ch/albert/alp-phd-soc-num-1-4-5.

(will be made public after acceptance of the manuscript)

## Code availability

All codes, executables, scripts, models, stimuli to reproduce the findings are available at https://gitlab.epfl.ch/albert/alp-phd-soc-num-1-4-5.

(will be made public after acceptance of the manuscript)

## Author contributions

L.A., F.B., B.H. and O.B designed study 1. L.A., J.P., F.B. and O.B designed study 2. L.A. realized the technological developments of study 1 and study 2. F.B., B.H, O.B. supervised study 1. F.B., J.P., O.B. supervised study 2. L.A. recruited participants and performed the experiments in study 1. L.A. analyzed the data in study 1 and study 2. L.A. and J.P recruited patients in study 2. J. P coordinated patient recruitment via patient associations: Parkinson Schweiz, Parkinson’s UK and Association France Parkinson. J.P. designed the questionnaire for the assessment of hallucinations in study 2. L.A., F.B. and O.B. wrote the manuscript. All authors provided critical revisions and approved the final version of the manuscript.

## Acknowledgements

This research was supported by two generous donors advised by CARIGEST SA (Fondazione Teofilo Rossi di Montelera e di Premuda and a second one wishing to remain anonymous) to O.B.; Bertarelli Novartis Foundation for Medical-Biological Research Foundation to O.B. The funder played no role in study design, data collection, analysis and interpretation of data, or the writing of this manuscript.

The authors thank all patients for their participation in study 2, as well as Parkinson Schweiz, Parkinson’s UK and Association France Parkinson in their help of recruiting patients with PD for study 2. The authors thank all participants for their participation in study 1. The authors thank Mr. Laurent Jenni for his robotics support. The authors thank Mr. Arthur Trivier for his help in the design of the robotic system figure part used in Figure 1 (modified from (Bernasconi et al. 2021)) and his help in the design of the desk figure part used in Figure 5.

## Competing interests

A patent application has been submitted based on the methods and results (Methods and systems to measure and quantify cognitive impairment in numerosity estimation) with L.A., O.B., F.B., B.H., and J.P. as inventors. O.B. is inventor on patent US 10,286,555 B2 held by the Swiss Federal Institute (EPFL) that covers the robot-controlled induction of presence hallucination. O.B. is inventor on patent US 10,349,899 B2 held by the Swiss Federal Institute (EPFL) that covers a robotic system for the prediction of hallucinations for diagnostic and therapeutic purposes. O.B. is co-founder and shareholder of Metaphysiks Engineering SA, a company that develops immersive technologies, including applications of the robotic induction of presence hallucinations that are not related to the diagnosis, prognosis or treatment of Parkinson’s disease. O.B. is a member of the board and shareholder of Mindmaze SA.

## Extended Data

**Extended Data Figure 1.**
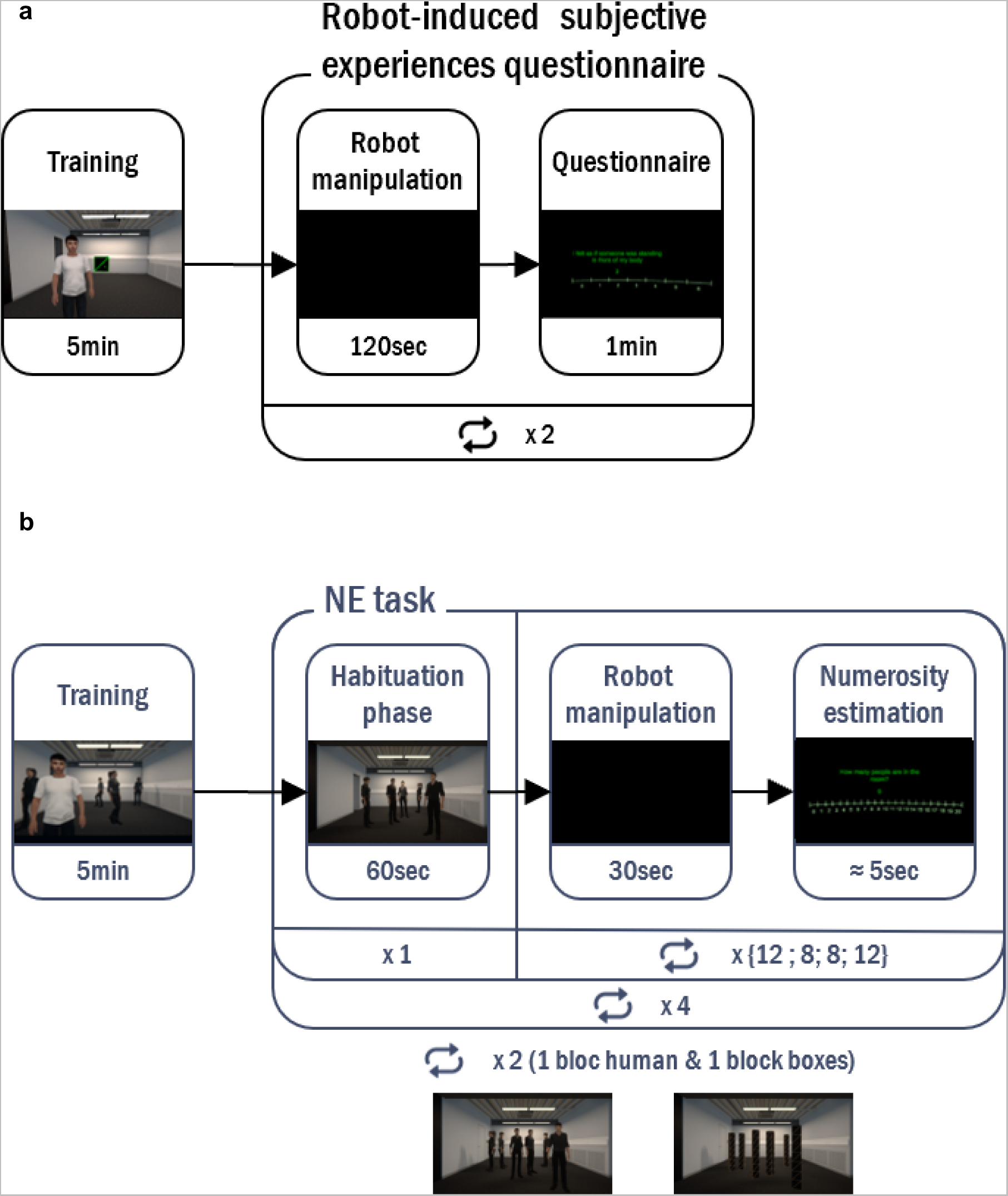
Experimental protocol (study 1). Schematic representation of the experimental protocol of the a) robot-induced subjective experiences questionnaire and b) numerosity estimation task (study 1). A detailed view of a numerosity trial (robot manipulation followed by numerosity estimation) is depicted in Figure 1.

