## Supplementary material for "Digital-robotic markers for hallucinations in Parkinson’s disease"

Olaf Blanke

Bertarelli Chair in Cognitive Neuroprosthetics

Neuro-X Institute

School of Life Sciences

Campus Biotech

Swiss Federal Institute of Technology

Ecole Polytechnique Fédérale de Lausanne (EPFL)

CH – 1012 Geneva



##### Keywords

Neurodegeneration, Virtual Reality, Robotics, Web-based assessment, Psychosis

#### Supplementary note 1: NEH subitizing range calculation (online pilot study)

An online web-based experiment was developed to determine the subitizing range of our numerosity stimuli. Participants first filled in some socio-demographic information (gender and age), followed by our NEH task. This experiment was available in French and English.

##### Study population

Twenty-height healthy participants (16 women, 12 men; age ranging from 21 to 42 years, mean ± SD age = 29.5 ± 6.13 years) took part in the online web-based experiment.

##### Numerosity stimuli

Visual stimuli were generated on Unity 3D (version 2019.3.13f1). The virtual environment was modeled in 3DS max and consisted of a realistic representation of our experimental room. Stimuli consisted of a 3D scene of our virtual experiment room with digital humans inside (examples in Figure S1a, Figure S1c, Figure S1e, Figure S1g). Digital humans were placed in the virtual scene in front of the camera viewpoint, facing the viewpoint in a range from – 90° to 90°, and in a way that they do not overlap completely from the viewpoint. Digital humans were placed between 1.75m and 5.25m in depth from viewpoint, and between -1.5m and 1.5m from right to left. The array of digital humans occupied a maximum virtual camera visual angle of 60° horizontally. This ensures that digital humans and control objects were located within the participants' peripheral field of view, which is the "vision produced by light falling on areas of the retina outside the macula" (Morris, Press, and Morris 1992), and that they were displayed inside and close to the limit of 30° retinal eccentricity, which is the limit from which the visual acuity decreases more strongly (Anstis 1974). The number of digital humans ranged from 1 to 24.

#### Procedure

Participants first filled in some socio-demographic information (gender and age). Then, before starting the NEH task, participants performed a screen calibration procedure, which consisted in measuring and reporting the length of a line displayed on their screen. This measure allowed to scale stimuli, so they were displayed the same physical size on each participant screen, independent of monitor screen and positions. Participants were instructed to stay fifty centimeters away from their screen. The NEH task contained 120 trials, 5 trials per numerosity (ranging from 1 to 24), randomized order between participants. Before each trial, participants were asked to fix their gaze on a fixation cross, drawn from a uniform distribution between 700 and 1500 msec (step of 200 msec). The total duration of the experiment was approximately 10minutes.

#### Data analysis

The error rate (ER), mean response time (RT), mean response, and variation coefficient (VC; standard deviation of response divided by mean response) were calculated for each numerosity and each participant. A stable VC across numerosity reflects scalar variability and Weber's law, which is a signature of estimation processes.

The proxy for the subitizing range was calculated using the ISR algorithm (Leibovich-Raveh et al. 2018). The data for RT (numerosity 1 to 7 (Revkin et al. 2008)) were used to fit a sigmoid function with unknown parameters using the Levenberg-Marquardt non-linear fitting algorithm (Marquardt 1963). The proxy for the subitizing range was taken as the intersection point between the tangent at the inflection point of the sigmoid curve and a line with a slope of zero and an intercept where the sigmoid curve crosses the y-axis at $x = 0$ (i.e., the subitizing line). This represents the point at which the slope begins to appreciably change, thus representing an accurate estimate of the upper bound of the subitizing range (Leibovich-Raveh et al. 2018). Linear mixed effects models (packages lme4 (Bates et al. 2015) and lmerTest (Kuznetsova, Brockhoff, and Christensen 2017) in R (Team 2013) with numerosity as a fixed effect and participant as a random effect were then used to examine the limit when participants' patterns of performance obeyed Weber's law by showing scalar variability (i.e., constant coefficients of variation), to confirm when estimation mechanisms were active.

##### *Preprocessing*

For each participant, trials in which response time were superior to the mean plus three standard deviations were excluded, resulting in the exclusion of 1.55% of the trials across conditions and participants (52 trials over 3360). Trials in which participants answer contains three digits were excluded, as well as trials in which the participants answer contains two digits when the presented numerosity was in the subitizing range, resulting in the exclusion of 0.12% of the trials (4 trials over 3300).

#### Results

##### *Descriptive statistics*

To demonstrate the expected pattern of numerosity task, we plotted RT, ER and VC as a function of numerosity (figure Note 1). Errors were very rare for numerosity 1 to 4 and then the error rate steeply increased with numerosity. RT gradually increased from numerosity 1 to 8 with a slope that became steeper with numerosity between numerosity 1 to 5, and with a slope that then became less steep with numerosity between numerosity 5 to 8. After numerosity 8, RT reaches a kind of plateau. This sigmoidal RT curve behavior is typically observed in numerosity experiment (Kaufman et al. 1949).

##### *ISR calculation*

We calculated ISR according to RT for correct responses only. Three participants had $R^2$ value of 0.7 or less, which were considered unreliable (Leibovich-Raveh et al. 2018) and they were excluded from further analysis. Mean $R^2$ value for the rest of the participants was 0.88, with a standard deviation of 0.08. This analysis yielded a mean subitizing range of 3.31 (SD = 1.19).

##### *Scalar variability*

We ran a series of linear mixed effects models on coefficient of variation data with numerosity as a fixed effect and participant as a random effect. With data considering numerosity 1 to 24 (respectively 2 to 24, 3 to 24 and 4 to 24), the difference was statistically different ($F(1,642)=155.97$, $p<0.001$), respectively $F(1,614)=96.89$, $p<0.001$), $F(1,5862)=47.27$, $p<0.001$) and $F(1,558)=16.18$, $p<0.001$)). With data considering numerosity 5 and above to 24, the difference was not significant anymore.

##### *Lower bound of the estimation range*

The limit when participants' patterns of performance obeyed Weber's law by showing scalar variability (i.e., constant coefficients of variation) was numerosity 5. This result, coupled with the proxy result of subitizing range calculated with the ISR gave us the lower bound of the estimation range at numerosity 5.

#### Apparatus and material

This online experiment was developed in house using javascript (on client side: javascript, html, css; on server side: https server in node.js, nginx as a reverse proxy, running in two docker containers) and was hosted on an EPFL Server in a dedicated Virtual Machine (1xvCPU, 1GB RAM, 40GB HDD) in Demilitarized Zone. Data were saved and stored on EPFL’s servers.

#### Ethics

This online study, as being irreversibly anonymous, was considered as falling outside of the scope of the Swiss legislation regulating research on human subjects, so that the need for local ethics committee approval was waived.

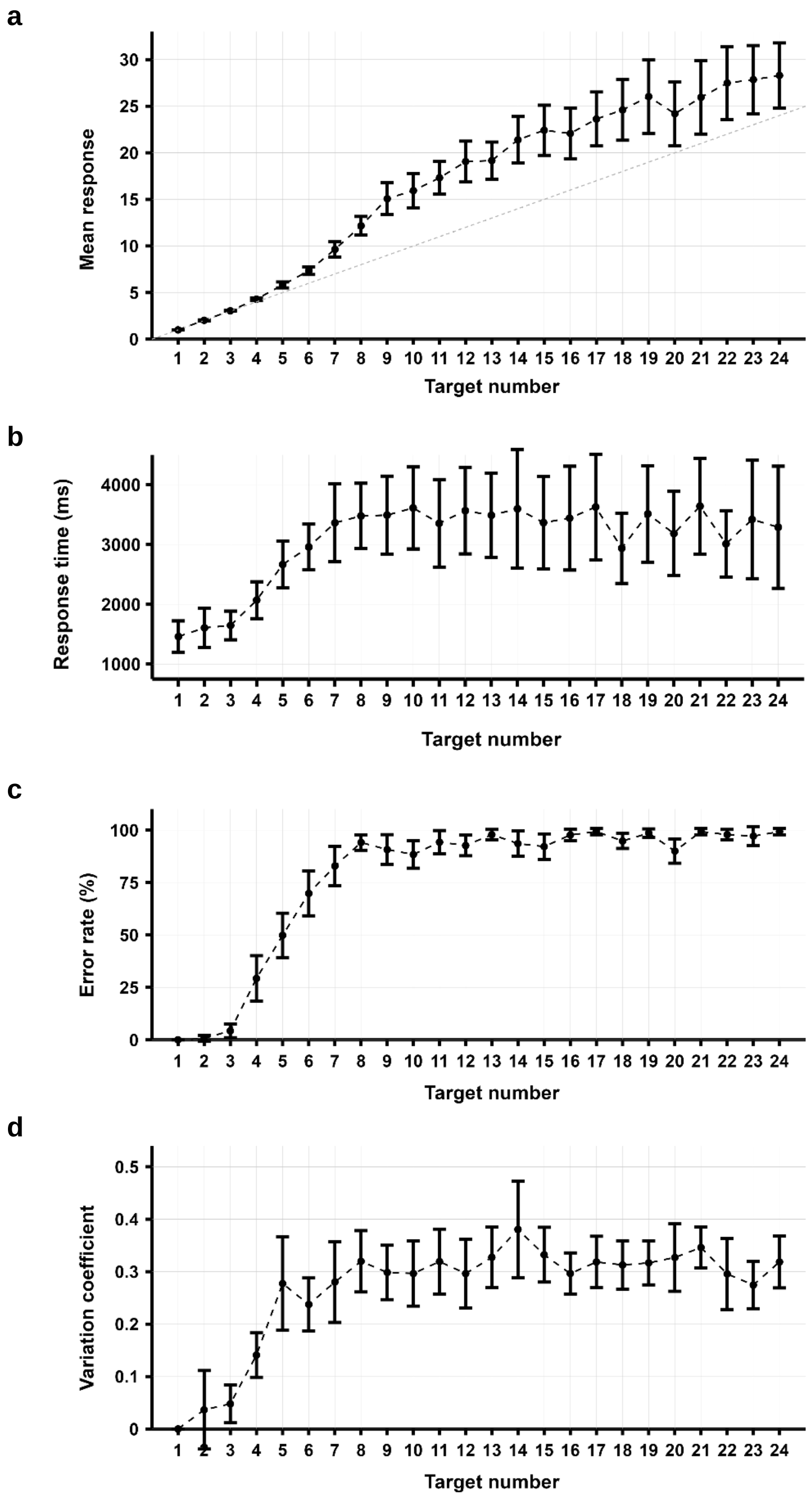


**Figure Note 1. Online pilot study results.** Subitizing range calculation experiment – Results. Mean response (a), response time (b), error rate (c) and variation coefficient (d) as a function of presented numerosity. Error bars represent 95 CI.

### Supplementary note 2: Mediation analysis NEH task (study 1)

#### Step 1: effect of the robotic sensorimotor stimulation on NEH

In step one of the mediation model, the effect of the robotic sensorimotor stimulation on NEH, ignoring the mediator (robot-induced PH question rating), was significant ($F(1,27)=26.05$; $p<0.001$).

| | Sum Sq | Mean Sq | NumDF | DenDF | F value | Pr(>F) |
|---|---|---|---|---|---|---|
| ***Robotic Sensorimotor Stimulation*** | 0.31 | 0.31 | 1 | 27 | 26.05 | <0.001 |

| | NEH | | |
|---|---|---|---|
| **Predictors** | **Estimates** | **CI** | **p** |
| **(Intercept)** | 6.82 | 6.62 – 7.20 | <0.001 |
| ***Robotic Sensorimotor Stimulation*** | 0.15 | 0.09 – 0.21 | <0.001 |
| **Random effects** | | | |
| **σ2** | 0.01 | | |
| ***τ00 S_ID*** | 0.27 | | |
| ***ICC*** | 0.96 | | |
| ***N S_ID*** | 28 | | |
| ***Observations*** | 56 | | |
| ***Marginal R2 / Conditional R2*** | 0.019/0.959 | | |

#### Step 2: effect of the robotic sensorimotor stimulation on robot-induced PH question rating

Step two showed that the effect of the robotic sensorimotor stimulation on the mediator (robot-induced PH question rating), was significant ($F(1,27)=10.44$; $p=0.003$).

| | Sum Sq | Mean Sq | NumDF | DenDF | F value | Pr(>F) |
|---|---|---|---|---|---|---|
| ***Robotic Sensorimotor Stimulation*** | 8.64 | 8.64 | 1 | 27 | 10.44 | 0.003 |

| | robot-induced PH question rating | | |
|---|---|---|---|
| **Predictors** | **Estimates** | **CI** | **p** |
| **(Intercept)** | 1.5 | 0.75 – 2.25 | <0.001 |
| ***Robotic Sensorimotor Stimulation*** | 0.79 | 0.30 – 1.27 | 0.002 |
| **Random effects** | | | |
| **σ2** | 0.83 | | |
| ***τ00 S_ID*** | 3.07 | | |
| ***ICC*** | 0.79 | | |
| ***N S_ID*** | 28 | | |
| ***Observations*** | 56 | | |
| ***Marginal R2 / Conditional R2*** | 0.039/0.796 | | |

#### Step 3: effect of the robot-induced PH question rating on NEH, controlling for robotic sensorimotor stimulation

Step three of the mediation process showed that the effect of the mediator (robot-induced PH question rating), controlling for robotic sensorimotor stimulation, was significant ($F(1,35)=6.53$; $p=0.02$). The effect of the robotic sensorimotor stimulation was also still significant ($F(1,28)=11.80$; $p=0.002$).

| | Sum Sq | Mean Sq | NumDF | DenDF | F value | Pr(>F) |
|---|---|---|---|---|---|---|
| ***Robotic Sensorimotor Stimulation*** | 0.12 | 0.12 | 1 | 28.13 | 11.80 | 0.002 |
| ***robot-induced PH question rating*** | 0.07 | 0.07 | 1 | 34.89 | 6.53 | 0.015 |

| | NEH | | |
|---|---|---|---|
| **Predictors** | **Estimates** | **CI** | **p** |
| **(Intercept)** | 6.75 | 6.54 – 6.95 | <0.001 |
| ***Robotic Sensorimotor Stimulation*** | 0.11 | 0.04 – 0.17 | 0.001 |
| ***robot-induced PH question rating*** | 0.05 | 0.01 – 0.09 | 0.014 |
| **Random effects** | | | |
| ***σ2*** | 0.01 | | |
| ***τ00 S_ID*** | 0.25 | | |
| ***ICC*** | 0.96 | | |
| ***N S_ID*** | 28 | | |
| ***Observations*** | 56 | | |
| ***Marginal R2 / Conditional R2*** | 0.056/0.962 | | |

#### Step 4: causal mediation analysis

ACME stands for average causal mediation effects (indirect effect of the robotic sensorimotor stimulation on the NEH), ADE stands for average direct effects (direct effect of the robotic sensorimotor stimulation on the NEH), Total Effect stands for the total effect (direct plus indirect effect of the robotic sensorimotor stimulation onto the NEH), Prop. Mediated describes the proportion of the effect of the robotic sensorimotor stimulation on the NEH that goes through the robot-induced PH question rating.

| | Estimates | 95% CI lower | 95% CI upper | p-value |
|---|---|---|---|---|
| **ACME** | 0.04 | 0.006 | 0.08 | 0.01 |
| ***ADE Stimulation*** | 0.11 | 0.05 | 0.17 | <0.001 |
| ***Total Effect*** | 0.15 | 0.09 | 0.21 | <0.001 |
| ***Pop. mediated*** | 0.26 | 0.04 | 0.56 | 0.01 |

#### Supplementary note 3: Mediation analysis NEO task (study 1)

##### Step 1: effect of the robotic sensorimotor stimulation on NEO

In step one of the mediation model, the effect of the robotic sensorimotor stimulation on NEO, ignoring the mediator (robot-induced PH question rating), was not significant ($F(1,27)=3.71$; $p=0.06$).

| | Sum Sq | Mean Sq | NumDF | DenDF | F value | Pr(>F) |
|---|---|---|---|---|---|---|
| ***Robotic Sensorimotor Stimulation*** | 0.13 | 0.13 | 1 | 27 | 3.71 | 0.065 |

| | NEO | | |
|---|---|---|---|
| **Predictors** | **Estimates** | **CI** | **p** |
| **(Intercept)** | 6.98 | 6.78 – 7.18 | <0.001 |
| ***Robotic Sensorimotor Stimulation*** | -0.09 | -0.19 – 0.00 | 0.06 |
| **Random effects** | | | |
| **σ2** | 0.03 | | |
| ***τ00 S_ID*** | 0.25 | | |
| ***ICC*** | 0.88 | | |
| ***N S_ID*** | 28 | | |
| ***Observations*** | 56 | | |
| ***Marginal R2 / Conditional R2*** | 0.008/0.883 | | |

##### Step 2: effect of the robotic sensorimotor stimulation on robot-induced PH question rating

Step two showed that the effect of the robotic sensorimotor stimulation on the mediator (robot-induced PH question rating), was significant ($F(1,27)=10.44$;$p=0.003$).

| | Sum Sq | Mean Sq | NumDF | DenDF | F value | Pr(>F) |
|---|---|---|---|---|---|---|
| ***Robotic Sensorimotor Stimulation*** | 8.64 | 8.64 | 1 | 27 | 10.44 | 0.003 |

| | robot-induced PH question rating | | |
|---|---|---|---|
| **Predictors** | **Estimates** | **CI** | **p** |
| **(Intercept)** | 1.5 | 0.75 – 2.25 | <0.001 |
| ***Robotic Sensorimotor Stimulation*** | 0.79 | 0.30 – 1.27 | 0.002 |
| **Random effects** | | | |
| **σ2** | 0.83 | | |
| ***τ00 S_ID*** | 3.07 | | |
| ***ICC*** | 0.79 | | |
| ***N S_ID*** | 28 | | |
| ***Observations*** | 56 | | |
| ***Marginal R2 / Conditional R2*** | 0.039/0.796 | | |

#### Step 3: effect of the robot-induced PH question rating on NEO, controlling for robotic sensorimotor stimulation

Step three of the mediation process showed that the effect of the mediator (robot-induced PH question rating), controlling for robotic sensorimotor stimulation, was not significant ($F(1,47)=0.008$; $p=0.93$). The effect of the robotic sensorimotor stimulation was also not significant ($F(1,28)=2.78$; $p=0.11$).

| | Sum Sq | Mean Sq | NumDF | DenDF | F value | Pr(>F) |
|---|---|---|---|---|---|---|
| ***Robotic Sensorimotor Stimulation*** | 0.10 | 0.10 | 1 | 27.63 | 2.78 | 0.11 |
| ***robot-induced PH question rating*** | 0.0003 | 0.0003 | 1 | 47.24 | 0.008 | 0.93 |

| | NEO | | |
|---|---|---|---|
| **Predictors** | **Estimates** | **CI** | **p** |
| **(Intercept)** | 6.98 | 6.76 – 7.21 | <0.001 |
| ***Robotic Sensorimotor Stimulation*** | -0.309 | -0.20 – 0.02 | 0.10 |
| ***robot-induced PH question rating*** | -0.00 | -0.07 – 0.06 | 0.93 |
| **Random effects** | | | |
| ***σ2*** | 0.03 | | |
| ***τ00 S_ID*** | 0.26 | | |
| ***ICC*** | 0.88 | | |
| ***N S_ID*** | 28 | | |
| ***Observations*** | 56 | | |
| ***Marginal R2 / Conditional R2*** | 0.008/0.883 | | |

#### Step 4: causal mediation analysis

ACME stands for average causal mediation effects (indirect effect of the robotic sensorimotor stimulation on the NEO), ADE stands for average direct effects (direct effect of the robotic sensorimotor stimulation on the NEO), Total Effect stands for the total effect (direct plus indirect effect of the robotic sensorimotor stimulation onto the NEO), Prop. Mediated describes the proportion of the effect of the robotic sensorimotor stimulation on the NEO that goes through the robot-induced PH question rating.

| | Estimates | 95% CI lower | 95% CI upper | p-value |
|---|---|---|---|---|
| **ACME** | -0.002 | -0.06 | 0.05 | 0.92 |
| ***ADE Stimulation*** | -0.09 | -0.20 | 0.02 | 0.10 |
| ***Total Effect*** | -0.09 | -0.19 | 0.00 | 0.06 |
| ***Pop. mediated*** | 0.02 | -1.19 | 1.18 | 0.93 |

#### Supplementary note 4: Effects of clinical variables on PH in PD and NE (study 2)

To ensure that the observed differences between NEH and NEO could not be explained by a difference in demographic and clinical variables, we built a linear mixed effects model with online NE magnitude as the outcome variable, PD group (PD-nH and PD-PH), presented numerosity and type of stimulus (digital humans or control objects) as fixed effect, age, gender, disease duration and first affected side as covariate fixed effects, and a random intercept for each subject. Our results show no influence of these covariates on NE (age ($F(1, 110)=0.66$; $p=0.42$); gender ($F(1, 110)=2.30$; $p=0.13$); disease duration ($F(1, 110)=2.79$; $p=0.10$); first affected side ($F(1, 110)=0.95$; $p=0.42$)) (Table S10), strengthening our argumentation that our selective online NEH effect in PD patients is directly linked to experiencing PH .

#### Supplementary note 5: Prevalence of hallucinations (study 2)

Over our 170 patients, 63% (108/170) reported having experienced hallucinatory phenomena (passage hallucination, presence hallucination, visual illusion or complex visual hallucination). Although the prevalence of hallucinatory symptoms in PD varies remarkably through studies, evidence from previous works suggest that Minor Hallucinations (MH) may occur in up to 72% of patients(Chang and Fox 2016; Fénelon and Alves 2010; Pacchetti et al. 2005), with PH and passage hallucinations being the most frequent MH(Fénelon et al. 2011; Fénelon and Alves 2010; Pagonabarraga et al. 2016). Our results corroborate this evidence with 63% (107/170) of our PD patients having experienced MH. The most frequent type of MH was passage hallucination in 53% of patients (90/170), followed by PH in 41% of patients (69/170). The frequencies and occurrences of the different forms of hallucinations are shown in Figure S13 and reported in Table S8.

#### Supplementary note 6: Online NEH and online NEO response time differences (study 2)

Additional analysis ensured that the observed differences between online NEH and online NEO are not explained by differences in task difficulty between groups. Thus, a linear mixed effects model with response time as the outcome variable, PD group (PD-nH and PD-PH), presented numerosity and type of stimulus (humans or control objects) as fixed effect, and a random intercept for each subject (Table S11) showed no differences in response times between the two PD groups (i.e., PD group ($F(1,116) = 0.58$; $p=0.45$); no main effect nor any interaction), suggesting each task (NEH and NEO) difficulty is equal between our two PD groups (PD-nH and PD-PH). However, our results indicate a main effect of type of stimuli ($F(1,9084)=78.67$; $p<0.001$), with participants being faster on the online NEO compared to online NEH (Table S11). This main effect was not expected, and suggest that, for our PD patients' sample, the NEO task is easier compared to the NEH task. This is supported by the fact PD patients gives higher NEH than NEO (main effect of Type of Stimuli ($F(1,9085)=1188$; $p<0.001$) in the previous model with online NE magnitude as the outcome variable, see Table S9). Although unexpected, we argue that this difference alone cannot explain the overestimation observed in PD-PH versus PD-nH and in online NEH versus online NEO, which is more prone to be intrinsically linked to the experience of spontaneous PH. Indeed, our observations follows our findings from study 1, with PD-PH overestimating NEH as compared to PD-nH. As in study 1, we observed no differences in NEO between groups. Furthermore, our results indicating no difference in response time between our two PD groups, this suggests that despite the fact the NEO task is easier compared to the NEH task in our patient's sample, each task difficulty is equal between our two PD groups (PD-nH and PD-PH). Taken together, this suggests that the overestimation observed in PD-PH versus PD-nH and in online NEH versus online NEO cannot be explained by a difference in task difficulty but is intrinsically linked to the experience of spontaneous PH.

#### Supplementary note 7: PD-nH vs PD-PH technical characteristics (study 2)

##### No statistical difference (Calibration scale factor)

| | Levene's Test for Equality of Variances | | *t-test for Equality of Means* | | | | |
|---|---|---|---|---|---|---|---|
| | **F** | **P-value** | **t** | **df** | ***P-value (two-tailed)*** | **95% CI lower** | **95% CI upper** |
| ***Calibration scale factor*** | 0.087 | 0.77 | 0.83 | 115.53 | 0.41 | -0.058 | 0.14 |

##### No statistical difference (Refresh rate)

| | Levene's Test for Equality of Variances | | *t-test for Equality of Means* | | | | |
|---|---|---|---|---|---|---|---|
| | **F** | **P-value** | **t** | **df** | ***P-value (two-tailed)*** | **95% CI lower** | **95% CI upper** |
| ***Refresh rate*** | 0.85 | 0.36 | -1.21 | 105.44 | 0.23 | -8.79 | 2.13 |

##### No statistical difference (Resolution)

| | Levene's Test for Equality of Variances | | *t-test for Equality of Means* | | | | |
|---|---|---|---|---|---|---|---|
| | **F** | **P-value** | **t** | **df** | ***P-value (two-tailed)*** | **95% CI lower** | **95% CI upper** |
| ***Resolution*** | 1.12 | 0.29 | -0.96 | 96.47 | 0.34 | -477625 | 165383 |

##### No statistical difference (use of a computer or tablet)

Fisher's exact test for count data ($p = 0.85$)

| | *PD-nH* | *PD-PH* |
|---|---|---|
| ***Computer*** | 36 | 43 |
| ***Tablet*** | 19 | 20 |

**No statistical difference (OS family)**
Fisher's exact test for count data (p = 0.72)

| | *PD-nH* | *PD-PH* |
|---|---|---|
| ***Windows*** | 32 | 43 |
| ***Mac OSX*** | 11 | 11 |
| ***Android*** | 7 | 4 |
| ***IOS*** | 3 | 2 |
| ***Linus*** | 2 | 2 |
| ***Chrome OS*** | 0 | 1 |

**No statistical difference (Browser family)**
Fisher's exact test for count data (p = 0.40)

| | *PD-nH* | *PD-PH* |
|---|---|---|
| ***Chrome*** | 25 | 24 |
| ***Firefox*** | 10 | 8 |
| ***Safari*** | 9 | 11 |
| ***Edge*** | 5 | 14 |
| ***Mobile Safari*** | 2 | 2 |
| ***Samsung internet*** | 1 | 3 |
| ***Chrome mobile*** | 0 | 1 |
| ***Chrome mobile iOS*** | 1 | 0 |
| ***Facebook*** | 1 | 0 |
| ***Amazon Silk*** | 1 | 0 |

#### Supplementary note 8: online study dropout rate (study 2)

13 PD patients (13/183 = 7%) stopped the experiment during the NE task.

All drops were made during the first half of the NE task (trials where drop happened: 3; 10; 14; 17; 20; 20; 22; 22; 25; 32; 38; 40; 40 – median = 22).

Participants could stop the experiment whenever they wanted, and they could indicate a reason for the dropout by ticking a box.

| Drop reason | count |
|---:|---|
| **The experiment takes too much time** | 5 |
| **The task is too complicated** | 0 |
| **The experiment is boring** | 4 |
| **I am too tired to continue** | 1 |
| **I am not satisfied with my answers** | 0 |
| **Other (not indicated)** | 3 |

Supplementary table 1. Statistical results for the ratings of the robot-induced sensation questionnaire (study 1)

| Question | Synchronous Mean | SD | Asynchronous Mean | SD | $X^2$ value | df | N | Significance |
|---|---|---|---|---|---|---|---|---|
| ***I felt as if I was touching my back myself*** | 3.00 | 2.13 | 2.04 | 2.01 | 5.78 | 1 | 28 | 0.016 |
| ***I felt as if someone else's was touching my back*** | 2.25 | 2.01 | 3.00 | 2.13 | 10.40 | 1 | 28 | 0.001 |
| ***I felt as if someone was standing close to me*** | 1.50 | 1.99 | 2.29 | 1.96 | 12.00 | 1 | 28 | <0.001 |
| ***I felt as if I was not controlling my movements or actions*** | 1.14 | 1.53 | 2.61 | 2.10 | 11.20 | 1 | 28 | <0.001 |
| ***I felt as if someone was standing in front of me*** | 0.57 | 1.14 | 0.50 | 0.75 | 0.04 | 1 | 28 | 0.83 |
| ***I felt as if I had two body*** | 0.71 | 1.33 | 0.96 | 1.69 | 1.23 | 1 | 28 | 0.27 |
| ***I felt anxious/stressed*** | 0.86 | 1.30 | 1.18 | 1.49 | 2.19 | 1 | 28 | 0.14 |

#### Supplementary table 2. Statistical results for the general estimation performance (study 1).

One sample t-test between the presented numerosity and estimated numerosity for each stimuli type (digital humans or control objects) and each presented numerosity (A). Two-samples t-test for equality of means and Levene's test for equality of variance on estimated numerosity between stimuli type (digital humans or control objects) for each presented numerosity (B).

A)

| **Numerosity** | **Stimuli** | **Mean** | **SD** | **95% CI lower** | **95% CI upper** | **t value** | **df** | ***P-value (two-tailed)*** |
|---|---|---|---|---|---|---|---|---|
| ***5*** | Humans | 5.09 | 0.15 | 5.04 | 5.15 | 3.30 | 27 | 0.003 |
| | Boxes | 5.10 | 0.19 | 5.02 | 5.17 | 2.75 | 27 | 0.010 |
| ***6*** | Humans | 6.25 | 0.41 | 6.09 | 6.41 | 3.26 | 27 | 0.003 |
| | Boxes | 6.32 | 0.39 | 6.17 | 6.47 | 4.35 | 27 | <0.001 |
| ***7*** | Humans | 7.52 | 0.68 | 7.26 | 7.78 | 4.08 | 27 | <0.001 |
| | Boxes | 7.53 | 0.75 | 7.24 | 7.82 | 3.78 | 27 | <0.001 |
| ***8*** | Humans | 8.72 | 1.12 | 8.29 | 9.16 | 3.41 | 27 | <0.001 |
| | Boxes | 8.78 | 1.05 | 8.38 | 9.19 | 3.95 | 27 | <0.001 |

B)

| | **Levene's Test for Equality of Variances** | | ***t-test for Equality of Means*** | | | | |
|---|---|---|---|---|---|---|---|
| **Numerosity** | **F** | **P-value** | **t** | **df** | ***P-value (two-tailed)*** | **95% CI lower** | **95% CI upper** |
| ***5*** | 2.72 | 0.10 | -0.08 | 51.579 | 0.94 | -0.09 | 0.09 |
| ***6*** | 0.23 | 0.64 | -0.06 | 53.861 | 0.53 | -0.28 | 0.15 |
| ***7*** | 0.42 | 0.52 | -0.06 | 53.511 | 0.96 | -0.39 | 0.37 |
| ***8*** | 0.0003 | 0.99 | -0.21 | 53.78 | 0.83 | -0.64 | 0.52 |

#### Supplementary table 3: statistical results for the numerosity estimation task estimated numerosity (study 1)

ToS corresponds to the type of stimuli presented (digital humans or control objects). RSS corresponds to the robotic sensorimotor stimulation (synchronous or asynchronous). PN corresponds to the presented numerosity.

| | Sum Sq | Mean Sq | NumDF | DenDF | F value | Pr(>F) |
|---|---|---|---|---|---|---|
| ***ToS*** | 0.71 | 0.71 | 1 | 2197 | 1.00 | 0.32 |
| ***RSS*** | 0.40 | 0.40 | 1 | 2197 | 0.56 | 0.45 |
| ***PN*** | 4175.09 | 1391.70 | 3 | 2197 | 1945.81 | <0.001 |
| ***ToS:RSS*** | 8.26 | 8.26 | 1 | 2197 | 11.54 | <0.001 |
| ***ToS:PN*** | 0.46 | 0.15 | 3 | 2197 | 0.22 | 0.89 |
| ***RSS:PN*** | 1.19 | 0.40 | 3 | 2197 | 0.56 | 0.64 |
| ***ToS:RSS:PN*** | 1.78 | 0.59 | 3 | 2197 | 0.83 | 0.48 |

Estimated marginal means and contrasts of the interaction ToS:RSS by ToS. Values or RSS are either S corresponding to synchronous robotic sensorimotor stimulation or A corresponding to asynchronous robotic sensorimotor stimulation.

| ToS | RSS | emmean | SE | df | lower.CL | upper.CL |
|---|---|---|---|---|---|---|
| NEH | S | 6.82 | 0.10 | 33.24 | 6.59 | 7.05 |
| NEH | A | 6.97 | 0.10 | 33.24 | 6.74 | 7.20 |
| NEO | S | 6.98 | 0.10 | 33.24 | 6.75 | 7.21 |
| NEO | A | 6.89 | 0.10 | 33.24 | 6.65 | 7.12 |

| Contrast | ToS | estimate | SE | df | 95% CI lower | 95% CI upper | *t* | P-value |
|---|---|---|---|---|---|---|---|---|
| S – A | NEH | -0.15 | 0.05 | 2197 | -0.25 | -0.5 | -2.9 | 0.003 |
| S – A | NEO | 0.09 | 0.05 | 2197 | -0.004 | 0.19 | 1.9 | 0.06 |

#### Supplementary table 4: statistical results for the numerosity estimation task response time (study 1)

ToS corresponds to the type of stimuli presented (digital humans or control objects). RSS corresponds to the robotic sensorimotor stimulation (synchronous or asynchronous). PN corresponds to the presented numerosity.

| | Sum Sq | Mean Sq | NumDF | DenDF | F value | Pr(>F) |
|---|---|---|---|---|---|---|
| ***ToS*** | 1.32 | 1.32 | 1 | 2197 | 0.73 | 0.39 |
| ***RSS*** | 5.44 | 5.44 | 1 | 2197 | 3.01 | 0.08 |
| ***PN*** | 77.79 | 25.93 | 3 | 2197 | 0.14 | <0.001 |
| ***ToS:RSS*** | 0.001 | 0.001 | 1 | 2197 | 0.0008 | 0.98 |
| ***ToS:PN*** | 2.08 | 0.69 | 3 | 2197 | 0.38 | 0.76 |
| ***RSS:PN*** | 1.77 | 0.59 | 3 | 2197 | 0.33 | 0.81 |
| ***ToS:RSS:PN*** | 1.58 | 0.53 | 3 | 2197 | 0.29 | 0.83 |

#### Supplementary table 5. Statistical results for the general estimation performance (study 2).

One sample t-test between the presented numerosity and estimated numerosity for each stimuli type (digital humans or control objects) and each presented numerosity (A). Two-samples t-test for equality of means and Levene's test for equality of variance on estimated numerosity between stimuli type (digital humans or control objects) for each presented numerosity (B).

A)

| Numerosity | Condition | Mean | SD | 95% CI lower | 95% CI upper | t value | df | *P-value (two-tailed)* |
|---|---|---|---|---|---|---|---|---|
| ***5*** | Humans | 6.20 | 0.98 | 6.04 | 6.36 | 15.07 | 151 | <0.001 |
| | Boxes | 5.54 | 0.72 | 5.43 | 5.66 | 9.25 | 151 | <0.001 |
| ***6*** | Humans | 7.54 | 1.19 | 7.34 | 7.73 | 15.94 | 151 | <0.001 |
| | Boxes | 6.58 | 0.94 | 6.43 | 6.73 | 7.59 | 151 | <0.001 |
| ***7*** | Humans | 8.82 | 1.70 | 8.55 | 9.09 | 13.16 | 151 | <0.001 |
| | Boxes | 7.58 | 1.26 | 7.38 | 7.79 | 5.71 | 151 | <0.001 |
| ***8*** | Humans | 10.47 | 2.45 | 10.08 | 10.87 | 12.46 | 151 | <0.001 |
| | Boxes | 8.55 | 1.60 | 8.30 | 8.81 | 4.27 | 151 | <0.001 |

B)

| | **Levene's Test for Equality of Variances** | | ***t-test for Equality of Means*** | | | | |
|---|---|---|---|---|---|---|---|
| **Numerosity** | **F** | **P-value** | **t** | **df** | ***P-value (two-tailed)*** | **95% CI lower** | **95% CI upper** |
| ***5*** | 8.37 | 0.004 | 6.64 | 277.60 | <0.001 | 0.46 | 0.85 |
| ***6*** | 8.74 | 0.003 | 7.82 | 286.48 | <0.001 | 0.72 | 1.20 |
| ***7*** | 9.95 | 0.002 | 7.19 | 278.07 | <0.001 | 0.90 | 1.57 |
| ***8*** | 20.26 | <0.001 | 8.09 | 260.11 | <0.001 | 1.45 | 2.39 |

#### Supplementary table 6: self-report questionnaire on alteration of perception (study 2)

| *Participant ID* | *Visual disturbance* |
|---|---|
| *Passage hallucination* | Have you ever felt that someone, something or a shadow passing or moving on your side(s)? |
| *Presence hallucination* | Have you ever felt that someone is behind you or close to you when there is actually no one there? |
| *Visual illusion* | Have you ever seen something else instead of a real object or living thing? For example, seeing an animal instead of a bush, a stain on the floor transforms into a crawling insect, or stationary objects perceived as being in motion. |
| *Visual hallucination* | Have you ever seen objects, people, animals or scenes that others could not see and claim not to be real? |

#### Supplementary table 7: visual disturbances reported by PD patients that were rejected for the numerosity estimation task analysis (study 2)

| *Participant ID* | *Visual disturbance* |
|---|---|
| *pnj1_kxdax5e0-112a35c7-5746-66d1-cfdb-fa578961d8ca* | “Double vision, occasional hallucination” |
| *pnj1_ky2stwm4-88cc31f9-847e-61f1-ef9b-39b5de3f2251* | “yes . Things change colour and appearance” |
| *pnj1_kswwqp-eb784aee-b97b-2a9d-d051-16ff676a8585* | « mes paupilles deviennent lourdes et je n'arrive pas à garder les yeux ouverts et j'ai difficile à rouvrir les yeux » |
| *pnj1_kx67m1i3-1e0ff548-715f-c096-cecd-41e386659a9d* | “Yes, I see images of my children in places where they are not present. I see quick images running with the corner of my eyes. I see shoes walking on the floor » |
| *pnj1_kx8x13l7-bbfc5884-8005-c487-db78-9ad240b838da* | « Double vision » |
| *pnj1_l140oidg-727cee41-8f28-a9bc-2e29-8c2ec7677c8e* | « oui. Impression de flou, de voir à travers de l'eau » |
| *pnj1_l1g3lle8-b279f7f7-9ca7-c886-5dc7-7d2818cb46aa* | « voie double par moment » |
| *pnj1_l2kxn2kf-3356f4f5-5ffe-f74d-ea29-b23a689e4a65* | « diplopie » |
| *pnj1_l2puopo1-e92db4cb-614b-6f4c-95ca-5d56f98dd084* | « Troubles d'accommodation, diplopie, carré noir » |
| *pnj1_l3elk8ml-3a8cb639-341d-aec7-f1f9-45678f6b4734* | “shadows seen in peripheral vision” |

#### Supplementary table 8: Occurrences of the different types of experienced hallucinations (study 2)

Participants reported the occurrence as Never, Rarely (less than once a month), Occasionally (several times, but less than once a week), Frequently (several times a week, but less than once a day), or Daily (almost every day, several times a day)

A) Table with all participants (N=170)

| | ***Passage sensation*** | ***Presence hallucination*** | ***Visual Illusion*** | ***Visual hallucinations*** |
|---|---|---|---|---|
| ***Never*** | 80 | 101 | 110 | 138 |
| ***Rarely*** | 45 | 30 | 33 | 19 |
| ***Occasionally*** | 25 | 25 | 17 | 8 |
| ***Frequently*** | 16 | 10 | 4 | 5 |
| ***Daily*** | 4 | 4 | 6 | 0 |

A) Table with only the participants kept for numerosity estimation data analysis (N=118).

| | ***Passage sensation*** | ***Presence hallucination*** | ***Visual Illusion*** | ***Visual hallucinations*** |
|---|---|---|---|---|
| ***Never*** | 62 | 55 | 84 | 96 |
| ***Rarely*** | 20 | 26 | 18 | 13 |
| ***Occasionally*** | 19 | 23 | 10 | 6 |
| ***Frequently*** | 13 | 10 | 3 | 3 |
| ***Daily*** | 4 | 4 | 3 | 0 |

#### Supplementary table 9: statistical results for the numerosity estimation task estimated numerosity (study 2)

ToS corresponds to the type of stimuli presented (digital humans or control objects). PDG corresponds to the PD group (PD-nH or PD-PH). PN corresponds to the presented numerosity.

| | Sum Sq | Mean Sq | NumDF | DenDF | F value | Pr(>F) |
|---|---|---|---|---|---|---|
| ***ToS*** | 3140.26 | 3140.26 | 1 | 9084.91 | 1188.26 | <0.001 |
| ***PDG*** | 10.68 | 10.68 | 1 | 115.88 | 4.04 | 0.047 |
| ***PN*** | 16293.09 | 5431.03 | 3 | 9084.31 | 2055.08 | <0.001 |
| ***ToS:PDG*** | 142.22 | 142.22 | 1 | 9084.91 | 53.81 | <0.001 |
| ***ToS:PN*** | 524.67 | 174.89 | 3 | 9084.05 | 66.18 | <0.001 |
| ***PDG:PN*** | 132.50 | 44.17 | 3 | 9084.31 | 16.71 | <0.001 |
| ***ToS:RSS:PN*** | 41.86 | 13.95 | 3 | 9084.05 | 5.28 | 0.001 |

Estimated marginal means and contrasts of the interaction ToS:PDG by ToS.

| ToS | PDG | emmean | SE | df | lower.CL | upper.CL |
|---|---|---|---|---|---|---|
| NEH | PD-nH | 7.86 | 0.15 | 122.13 | 7.51 | 8.21 |
| NEH | PD-PH | 8.53 | 0.14 | 122.19 | 8.20 | 8.86 |
| NEO | PD-nH | 6.94 | 0.15 | 122.33 | 6.59 | 7.29 |
| NEO | PD-PH | 7.11 | 0.14 | 122.13 | 6.78 | 7.44 |

| Contrast | ToS | estimate | SE | df | 95% CI lower | 95% CI upper | *t* | P-value |
|---|---|---|---|---|---|---|---|---|
| PD-nH – PD-PH | NEH | -0.67 | 0.21 | 122.16 | -1.09 | -0.25 | -3.16 | 0.002 |
| PD-nH – PD-PH | NEO | -0.17 | 0.21 | 122.24 | -0.59 | 0.25 | -0.81 | 0.42 |

#### Supplementary table S10: statistical results for the numerosity estimation task estimated numerosity with covariates (study 2)

ToS corresponds to the type of stimuli presented (digital humans or control objects). PDG corresponds to the PD group (PD-nH or PD-PH). PN corresponds to the presented numerosity.

| | Sum Sq | Mean Sq | NumDF | DenDF | F value | Pr(>F) |
|---|---|---|---|---|---|---|
| ***ToS*** | 3140.53 | 3140.53 | 1 | 9084.96 | 1188.36 | <0.001 |
| ***PDG*** | 15.34 | 15.34 | 1 | 109.92 | 5.81 | 0.018 |
| ***PN*** | 16293.83 | 5431.28 | 3 | 9084.31 | 2055.17 | <0.001 |
| ***PD_side*** | 7.51 | 2.50 | 3 | 109.96 | 0.95 | 0.42 |
| ***PD_duration*** | 7.36 | 7.36 | 1 | 109.95 | 2.79 | 0.098 |
| ***age*** | 1.73 | 1.73 | 1 | 109.98 | 0.66 | 0.42 |
| ***gender*** | 6.08 | 6.08 | 1 | 109.92 | 2.30 | 0.13 |
| ***ToS:PDG*** | 142.12 | 142.12 | 1 | 9084.90 | 53.78 | <0.001 |
| ***ToS:PN*** | 524.57 | 174.86 | 3 | 9084.07 | 66.16 | <0.001 |
| ***PDG:PN*** | 132.41 | 44.14 | 3 | 9084.32 | 16.70 | <0.001 |
| ***ToS:RSS:PN*** | 41.87 | 13.96 | 3 | 9084.07 | 5.28 | 0.001 |

Estimated marginal means and contrasts of the interaction ToS:PDG by ToS.

| ToS | PDG | emmean | SE | df | lower.CL | upper.CL |
|---|---|---|---|---|---|---|
| NEH | PD-nH | 7.75 | 0.18 | 114.36 | 7.35 | 8.16 |
| NEH | PD-PH | 8.52 | 0.18 | 113.94 | 8.12 | 8.92 |
| NEO | PD-nH | 6.83 | 0.18 | 114.49 | 6.43 | 7.24 |
| NEO | PD-PH | 7.10 | 0.18 | 113.98 | 6.70 | 7.50 |

| Contrast | ToS | estimate | SE | df | 95% CI lower | 95% CI upper | *t* | P-value |
|---|---|---|---|---|---|---|---|---|
| PD-nH – PD-PH | NEH | -0.76 | 0.22 | 115.60 | -1.19 | -0.34 | -3.53 | <0.001 |
| PD-nH – PD-PH | NEO | -0.27 | 0.22 | 115.70 | -0.69 | 0.16 | -1.23 | 0.22 |

#### Supplementary table 11: statistical results for the numerosity estimation task response time (study 2)

ToS corresponds to the type of stimuli presented (digital humans or control objects). PDG corresponds to the PD group (PD-nH or PD-PH). PN corresponds to the presented numerosity.

| | Sum Sq | Mean Sq | NumDF | DenDF | F value | Pr(>F) |
|---|---|---|---|---|---|---|
| ***ToS*** | 199348501.1 | 199348501.1 | 1 | 9084.09 | 78.67 | <0.001 |
| ***PDG*** | 1463771.6 | 1463771.6 | 1 | 115.76 | 0.58 | 0.45 |
| ***PN*** | 379889853.3 | 126629951.1 | 3 | 9083.90 | 49.98 | <0.001 |
| ***ToS:PDG*** | 965158.8 | 965158.8 | 1 | 9084.09 | 0.38 | 0.54 |
| ***ToS:PN*** | 866669.4 | 288889.8 | 3 | 9083.81 | 0.11 | 0.95 |
| ***PDG:PN*** | 16896421.8 | 5632140.6 | 3 | 9083.90 | 2.22 | 0.08 |
| ***ToS:RSS:PN*** | 2195005.4 | 731668.5 | 3 | 9083.81 | 0.29 | 0.83 |

Estimated marginal means of ToS at each numerosity

| ToS | emmean | SE | df | lower.CL | upper.CL |
|---|---|---|---|---|---|
| NEH | 4617.16 | 181.23 | 117.97 | 4258.29 | 4976.04 |
| NEO | 4322.05 | 181.23 | 117.99 | 3963.15 | 4680.94 |

| Numerosity | Condition | Emmean | SE | df | 95% CI lower | 95% CI upper |
|---|---|---|---|---|---|---|
| ***5*** | Humans | 4295.843 | 185.7081 | 130.0746 | 3874.723 | 4716.963 |
| | Boxes | 4030.099 | 185.8071 | 130.3516 | 3608.765 | 4451.433 |
| ***6*** | Humans | 4589.319 | 185.7256 | 130.1235 | 4168.161 | 5010.476 |
| | Boxes | 4276.233 | 185.7261 | 130.1248 | 3855.074 | 4697.392 |
| ***7*** | Humans | 4720.750 | 185.7494 | 130.1900 | 4299.541 | 5141.959 |
| | Boxes | 4408.486 | 185.7673 | 130.2401 | 3987.238 | 4829.734 |
| ***8*** | Humans | 4862.735 | 185.7548 | 130.2053 | 4441.514 | 5283.956 |
| | Boxes | 4573.365 | 185.7606 | 130.2214 | 4152.131 | 4994.598 |

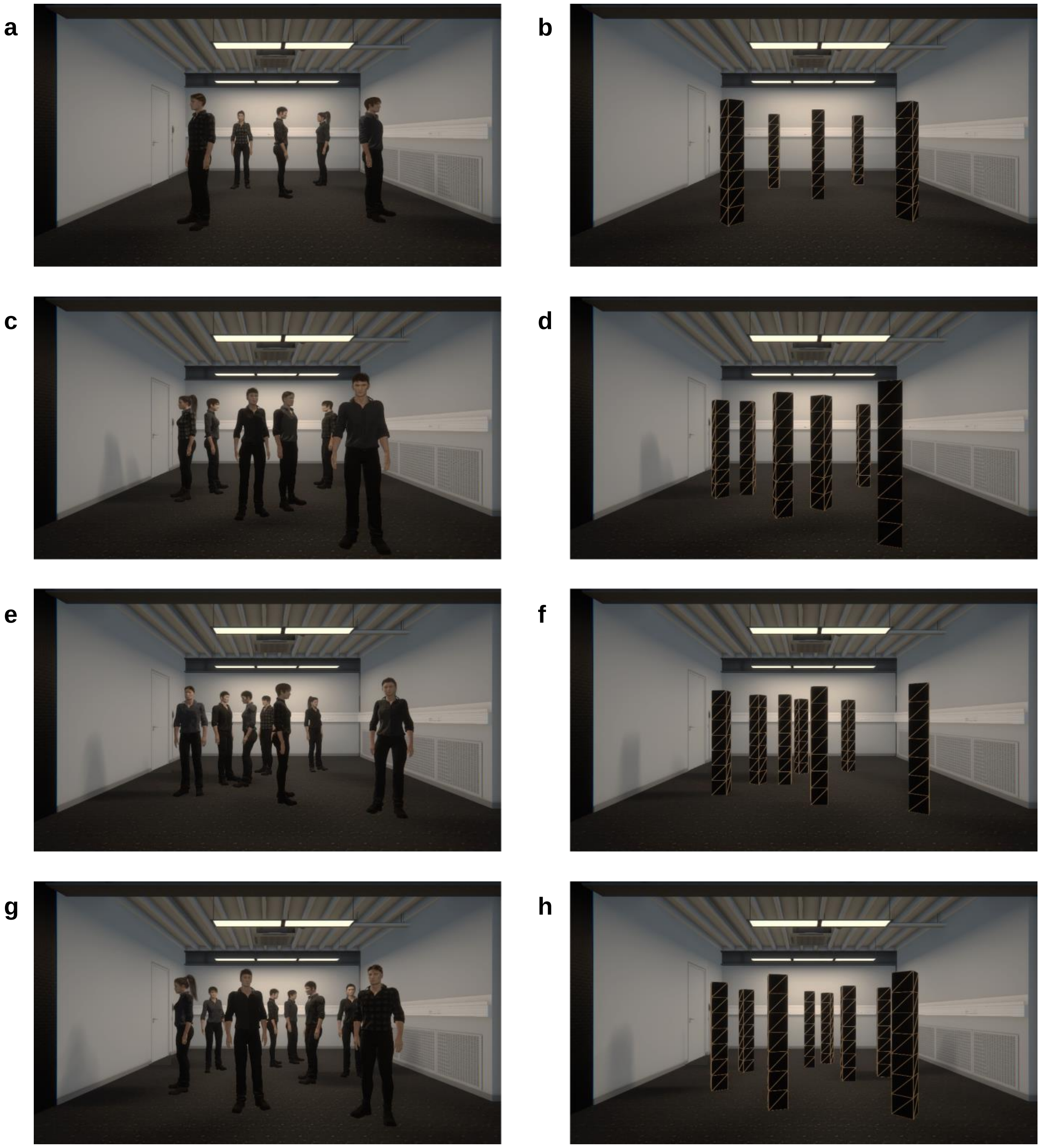


**Supplementary figure 1. Example of NEH and NEO stimuli.** Example with (a) five, (c) six, (e) seven, (g) eight humans (NEH) or (b) five, (d) six, (f) seven, (h) eight objects (NEO).

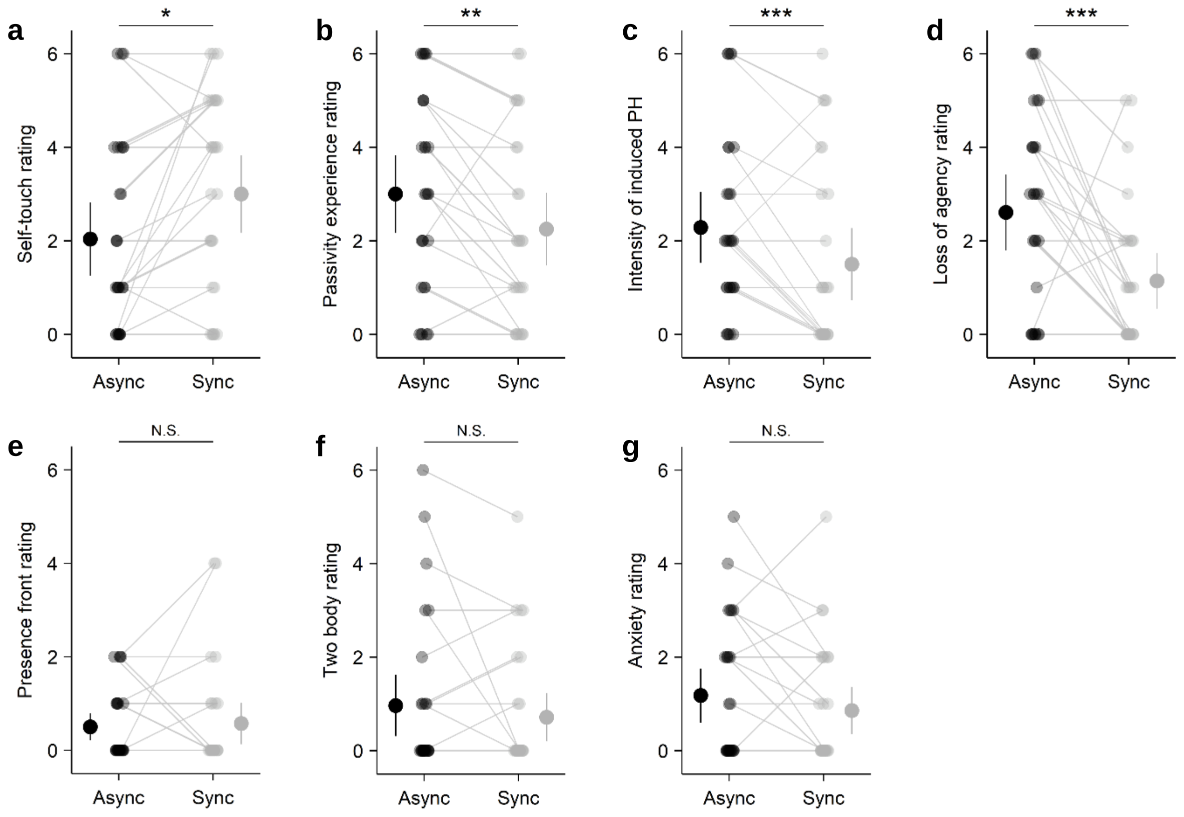


**Supplementary figure 2. Robot induced sensation questionnaire ratings (asynchronous versus synchronous) (study 1).** (a) Self-touch assessment ratings. (b) Passivity experience assessment ratings. (c) robot-induced PH assessment ratings. (d) Loss of agency assessment ratings. (e) Presence on the front assessment ratings. (f) Two-body assessment ratings. (g) Anxiety assessment ratings. Each linked pair of dots indicates the individual rating of the intensity of the assessed sensation (Asynchronous condition (dark grey) and synchronous condition (light grey)). The dots with the bar on the left and right sides indicate the mixed-effects linear regression between asynchronous (dark grey) and synchronous (light gray) sensorimotor stimulation. Error bar represents 95% confidence interval. *P ≤ 0.05 ; **P ≤ 0.01 ; ***P ≤ 0.001.

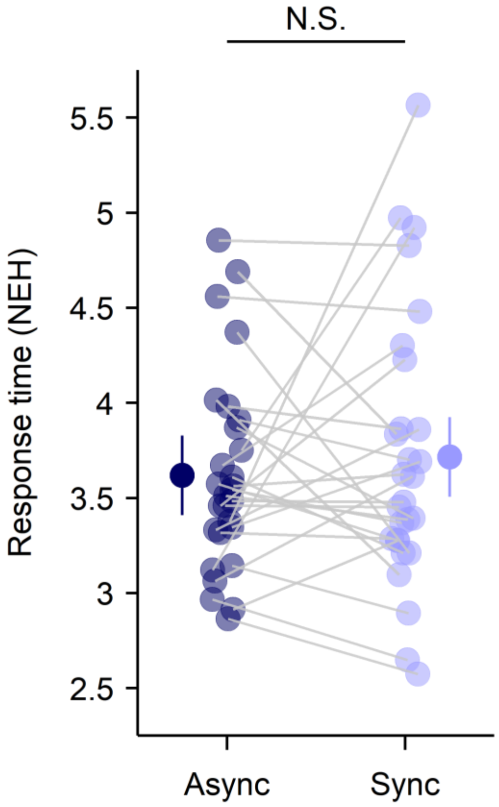


**Supplementary figure 3. NEH response time (synchronous, asynchronous) (study 1).** Each linked pair of dots indicates the individual NEH response time mean estimate in asynchronous condition (dark blue) and synchronous condition (light blue). The dots with the bar on the left and right sides indicate the mixed-effects linear regression between asynchronous (dark blue) and synchronous (light blue) sensorimotor stimulation. Error bar represents 95% confidence interval. N.S., not significant.

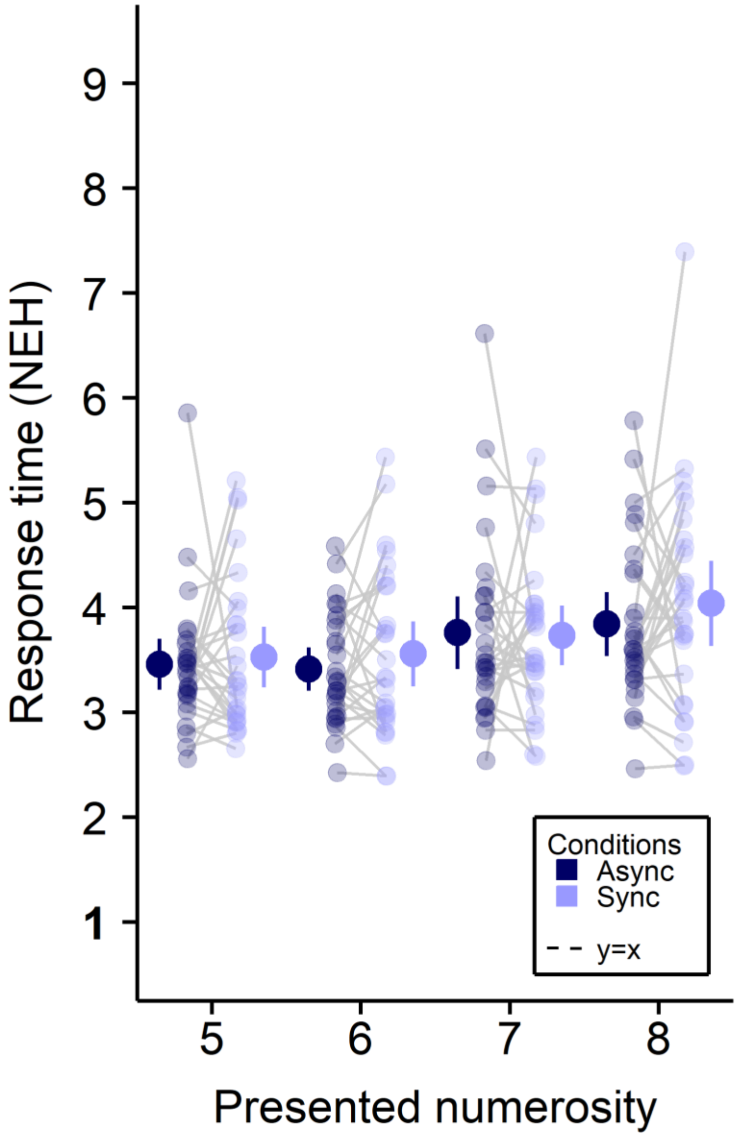


**Supplementary figure 4. NEH response time (synchronous, asynchronous) (study 1).** Task response time is shown for each presented numerosity in the NEH task, separately for the asynchronous and synchronous condition. Each linked pair of dots indicates the individual NEH response time mean estimate at each presented numerosity in asynchronous condition (dark blue) and synchronous condition (light blue). The dots with the bar on the left and right sides indicate the mixed-effects linear regression between asynchronous (dark blue) and synchronous (light blue) sensorimotor stimulation at each presented numerosity. Error bar represents 95% confidence interval.

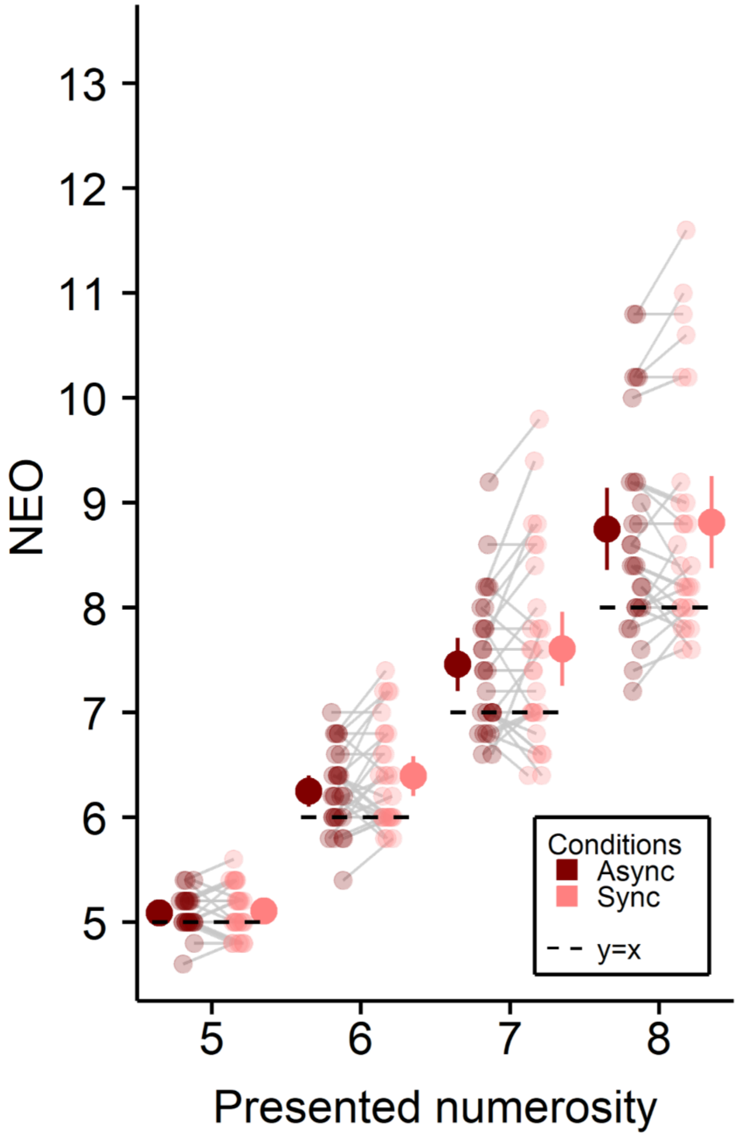


**Supplementary figure 5. NEO (each presented numerosity, synchronous, asynchronous) (study 1).** Task performance is shown for each presented numerosity in the NEO task, separately for the asynchronous and synchronous condition. Each linked pair of dots indicates the individual NEO mean estimate at the corresponding presented numerosity in asynchronous condition (dark red) and synchronous condition (light red). The dots with the bar on the left and right sides indicate the mixed-effects linear regression between asynchronous (dark red) and synchronous (light red) sensorimotor stimulation at each presented numerosity. Error bar represents 95% confidence interval.

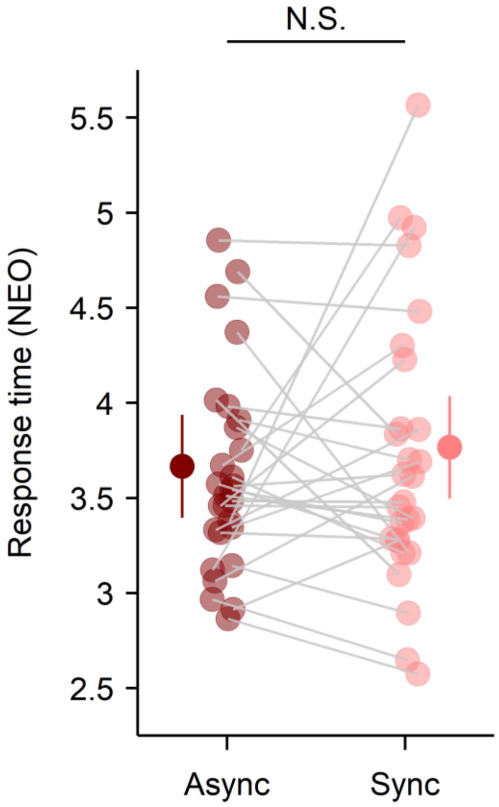


**Supplementary figure 6. NEO response time (synchronous, asynchronous) (study 1).** Each linked pair of dots indicates the individual NEO response time mean estimate in asynchronous condition (dark red) and synchronous condition (light red). The dots with the bar on the left and right sides indicate the mixed-effects linear regression between asynchronous (dark red) and synchronous (light red) sensorimotor stimulation. Error bar represents 95% confidence interval. N.S., not significant.

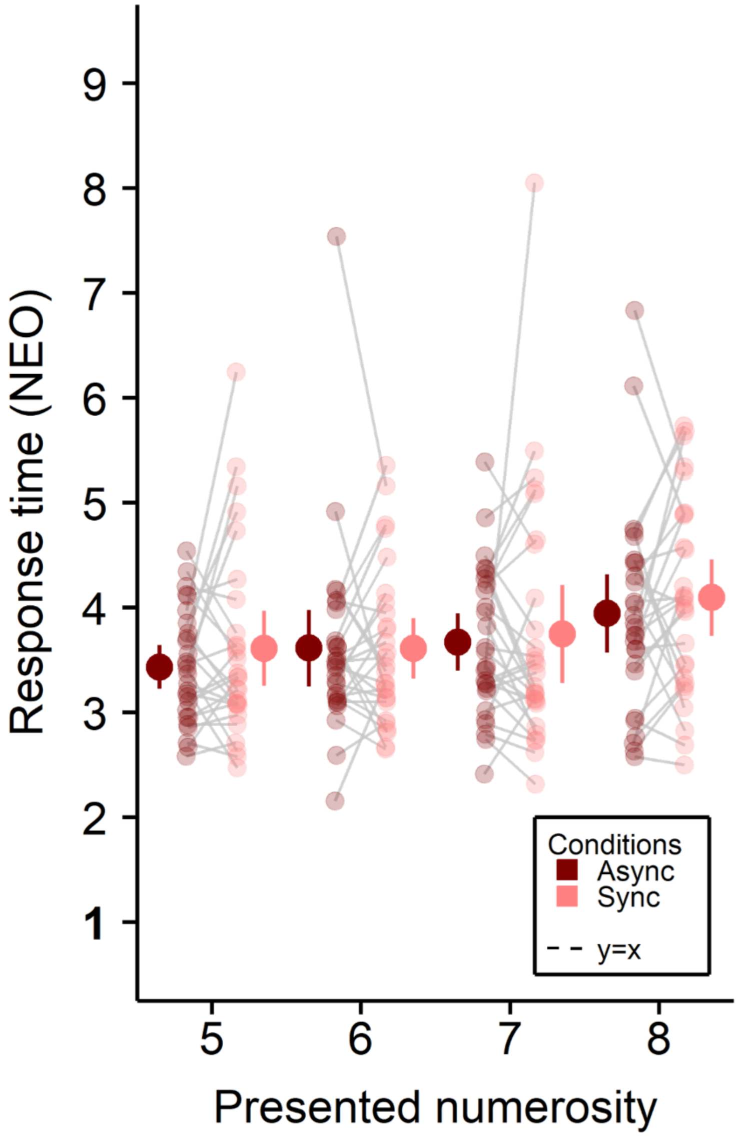


**Supplementary figure 7. NEO response time (synchronous, asynchronous) (study 1).** Task response time is shown for each presented numerosity in the NEO task, separately for the asynchronous and synchronous condition. Each linked pair of dots indicates the individual NEO response time mean estimate at each presented numerosity in asynchronous condition (dark red) and synchronous condition (light red). The dots with the bar on the left and right sides indicate the mixed-effects linear regression between asynchronous (dark red) and synchronous (light red) sensorimotor stimulation at each presented numerosity. Error bar represents 95% confidence interval.

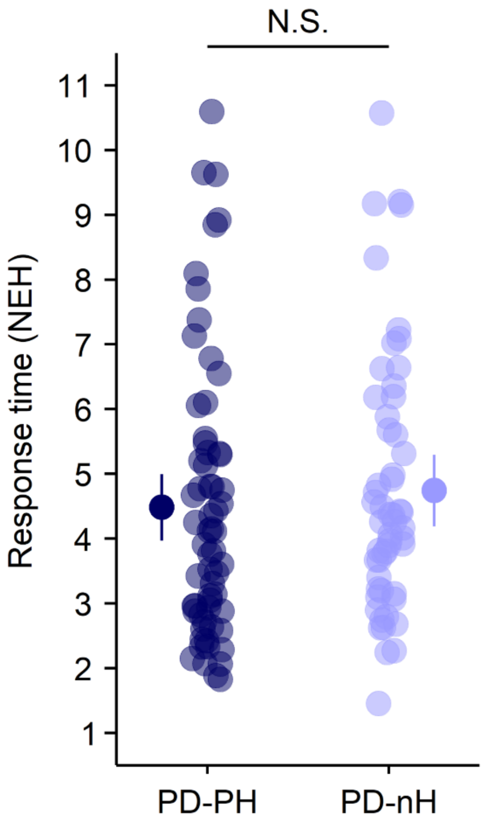


**Supplementary figure 8. NEH response time in PD patients (PD-PH and PD-nH) (study 2).** Each dot indicates the individual NEH response time mean estimate (PD-PH (dark blue) and PD-nH (light blue)). The dots with the bar on the left and right sides indicate the mixed-effects linear regression between PD-PH (dark blue) and PD-nH (light blue). Error bar represents 95% confidence interval. N.S., not significant.

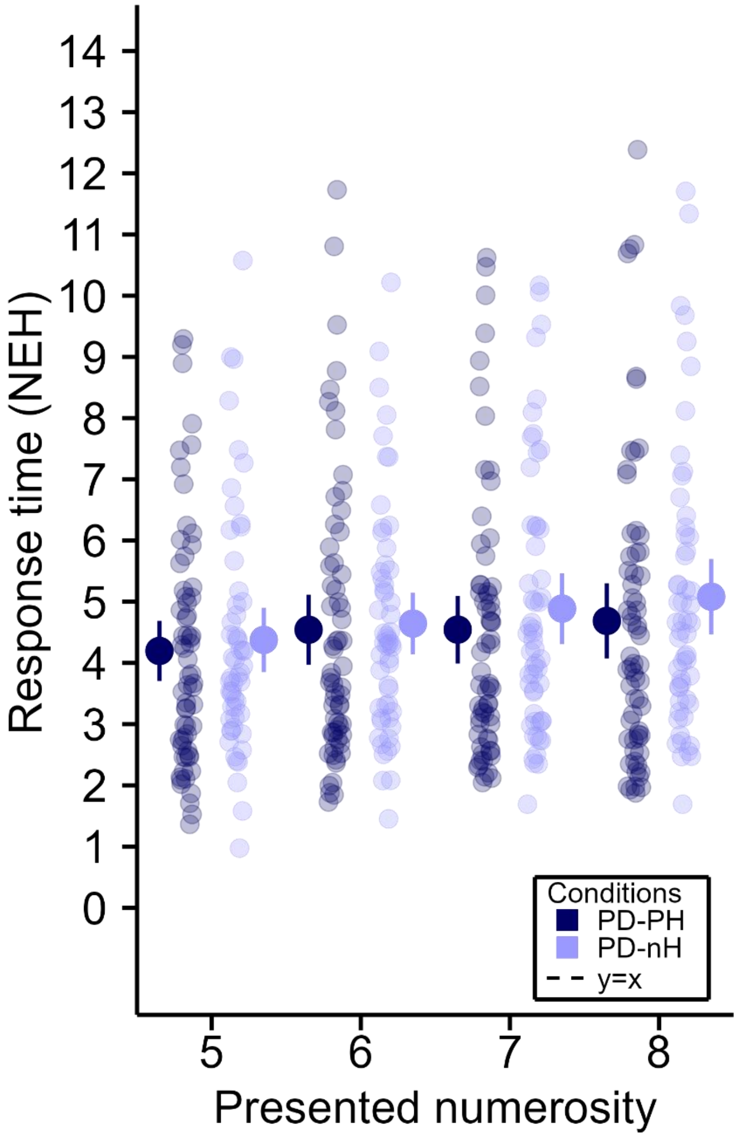


**Supplementary figure 9. NEH response time in PD patients (PD-PH and PD-nH) (study 2).** Response time is shown in PD patients for each tested numerosity in the NEH task for PD-PH and PD-nH separately. Each dot indicates the individual NEH response time mean estimate at the corresponding presented numerosity (PD-PH (dark blue) and PD-nH (light blue)). The dots with the bar on the left and right sides indicate the mixed-effects linear regression between PD-PH (dark blue) and PD-nH (light blue) at each presented numerosity. Error bar represents 95% confidence interval.

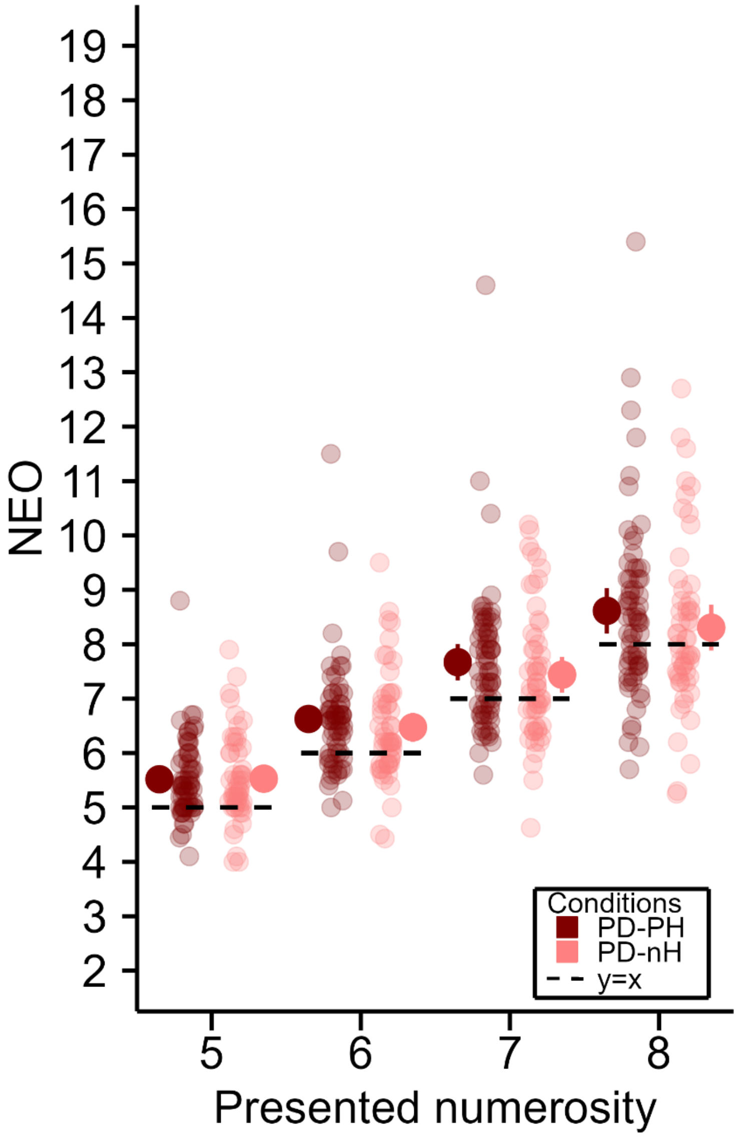


**Supplementary figure 10. NEO in PD patients (each presented numerosity, PD-PH vs PD-nH) (study 2).** Performance is shown in PD patients for each tested numerosity in the NEO task for PD-PH and PD-nH separately. Each dot indicates the individual NEO mean estimate at the corresponding presented numerosity (PD-PH (dark red) and PD-nH (light red)). The dots with the bar on the left and right sides indicate the mixed-effects linear regression between PD-PH (dark red) and PD-nH (light red) at each presented numerosity. Error bar represents 95% confidence interval.

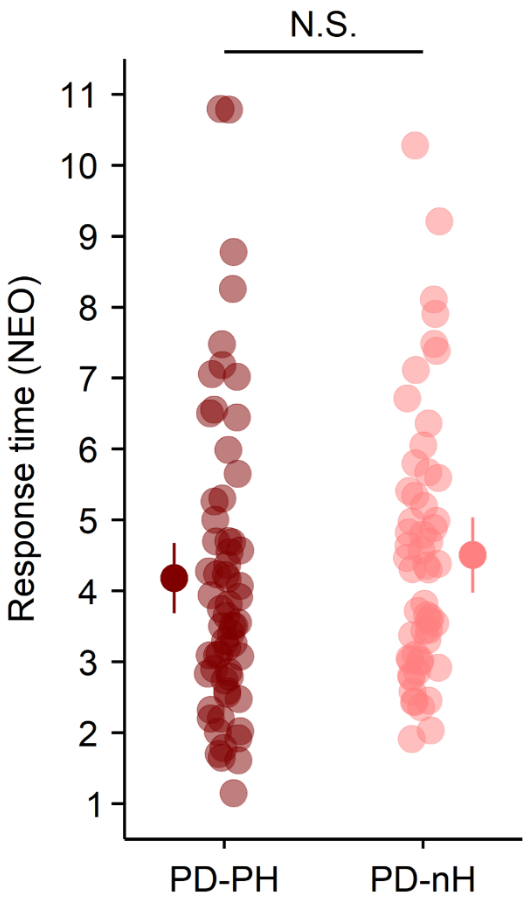


**Supplementary figure 11. NEO response time (asynchronous versus synchronous) (study 2).** Each linked pair of dots indicates the individual NEO response time mean estimate in asynchronous condition (dark red) and synchronous condition (light red). The dots with the bar on the left and right sides indicate the mixed-effects linear regression between asynchronous (dark red) and synchronous (light red) sensorimotor stimulation. Error bar represents 95% confidence interval. N.S., not significant.

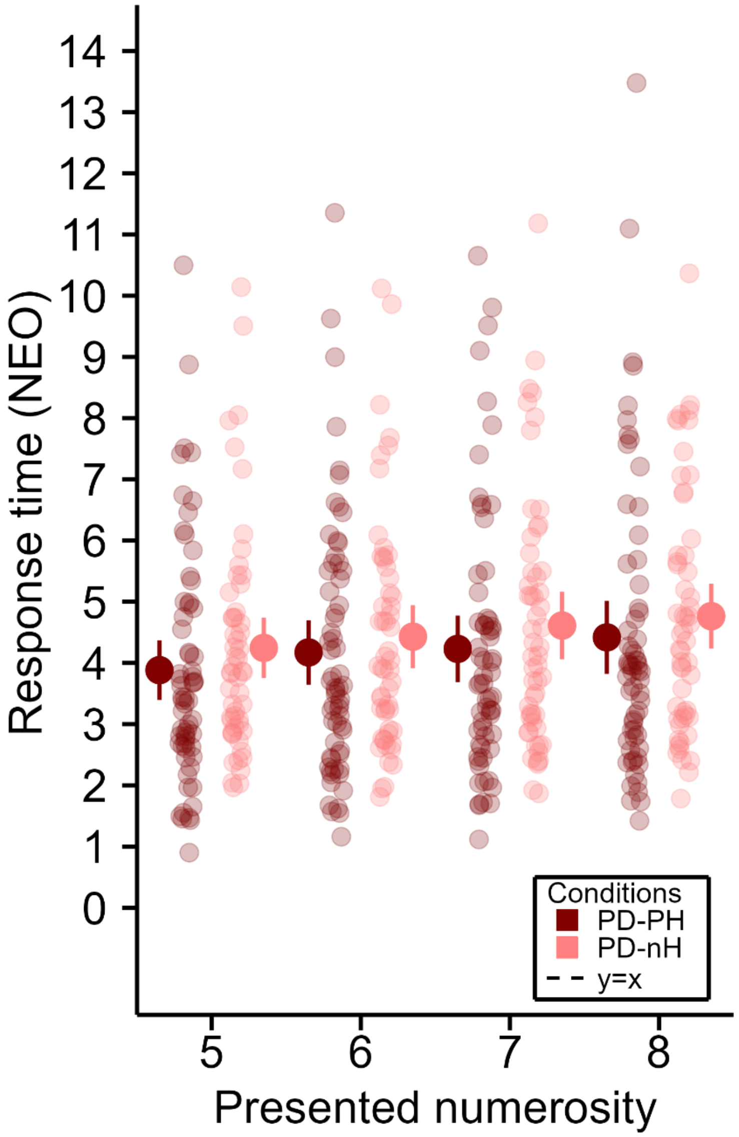


**Supplementary figure 12. NEO response time in PD patients (each presented numerosity, PD-PH vs PD-nH) (study 2).** Response time is shown in PD patients for each tested numerosity in the NEO task for PD-PH and PD-nH separately. Each dot indicates the individual NEO response time mean estimate at the corresponding presented numerosity (PD-PH (dark red) and PD-nH (light red)). The dots with the bar on the left and right sides indicate the mixed-effects linear regression between PD-PH (dark red) and PD-nH (light red) at each presented numerosity. Error bar represents 95% confidence interval.

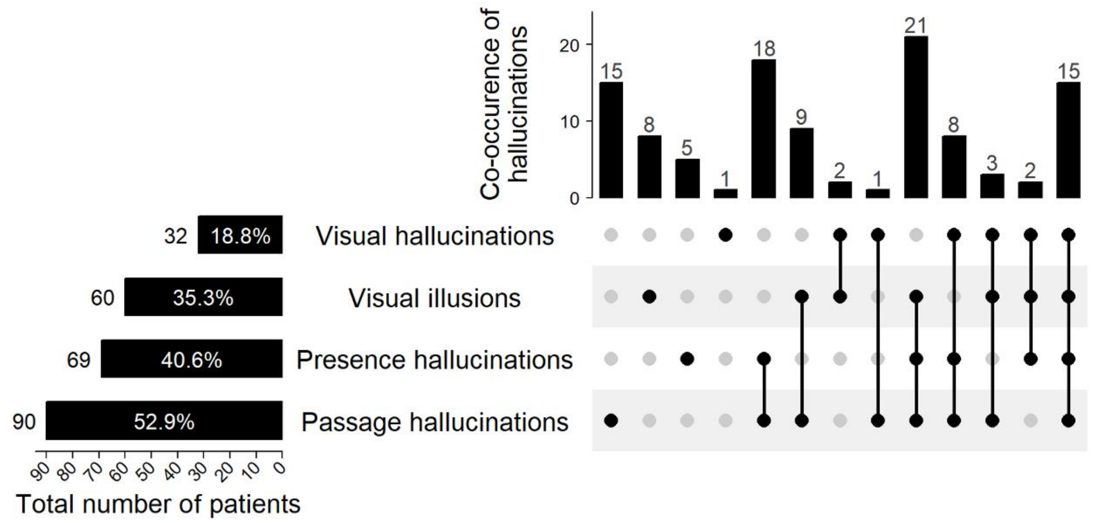


**Supplementary figure 13. Prevalence of hallucinations (study 2).** Prevalence of hallucination in PD (N=170). The sum of patients experiencing a specific hallucination is represented by the left bar plot. Every possible combination of multiple hallucination for a single patient is represented by the connected lines, and the number of patients is shown by the top bar plot.

Morris, Christopher G., Academic Press, and Christopher W. Morris. 1992. *Academic Press Dictionary of Science and Technology*. Gulf Professional Publishing.

Pacchetti, Claudio, Raffaele Manni, Roberta Zangaglia, Francesca Mancini, Enrico Marchioni, Cristina Tassorelli, Michele Terzaghi, et al. 2005. “Relationship between Hallucinations, Delusions, and Rapid Eye Movement Sleep Behavior Disorder in Parkinson’s Disease.” *Movement Disorders* 20 (11): 1439–48. https://doi.org/10.1002/mds.20582.

Pagonabarraga, Javier, Saul Martinez-Horta, Ramón Fernández de Bobadilla, Jesús Pérez, Roser Ribosa-Nogué, Juan Marín, Berta Pascual-Sedano, Carmen García, Alexandre Gironell, and Jaime Kulisevsky. 2016. “Minor Hallucinations Occur in Drug-Naive Parkinson’s Disease Patients, Even from the Premotor Phase.” *Movement Disorders* 31 (1): 45–52. https://doi.org/10.1002/mds.26432.

Revkin, Susannah K., Manuela Piazza, Véronique Izard, Laurent Cohen, and Stanislas Dehaene. 2008. “Does Subitizing Reflect Numerical Estimation?” *Psychological Science* 19 (6): 607–14. https://doi.org/10.1111/j.1467-9280.2008.02130.x.

Team, R. Core. 2013. “R: A Language and Environment for Statistical Computing.”
